# A common binding motif in the ET domain of BRD3 forms polymorphic structural interfaces with host and viral proteins

**DOI:** 10.1101/2020.09.21.306696

**Authors:** Sriram Aiyer, G.V.T. Swapna, Li-Chung Ma, Gaohua Liu, Jingzhou Hao, Gordon Chalmers, Brian C. Jacobs, Gaetano T. Montelione, Monica J. Roth

**Author notes:** Salk Institute for Biological Studies, 10010 North Torrey Pines Road, La Jolla, CA, 92037, USA. Nexomics Bioscience, Inc. 5 Crescent Ave, Rocky Hill, NJ 08553. Authors contributed equally to this work (Sriram Aiyer, G.V.T. Swapna).

## Abstract

The extra-terminal (ET) domain of BRD3 is conserved among BET proteins (BRD2, BRD3, BRD4), interacting with multiple host and viral protein-protein networks. Solution NMR structures of complexes formed between BRD3-ET domain with either the 79-residue murine leukemia virus integrase (IN) C-terminal domain (IN_329-408_), or its 22-residue IN tail peptide (TP) (IN_386-407_) alone, reveal similar intermolecular three-stranded β-sheet formation. ^15^N relaxation studies reveal a 10-residue linker region (IN_379-388_) tethering the SH3 domain (IN_329-378_) to the ET-binding motif (IN_389-405_)-ET complex. This linker has restricted flexibility, impacting its potential range of orientations in the IN - nucleosome complex. The complex of the ET-binding peptide of host NSD3 protein (NSD3_148-184_) and BRD3-ET domain includes a similar three-stranded β-sheet interaction, but the orientation of the β−hairpin is flipped compared to the two IN : ET complexes. These studies expand our understanding of molecular recognition polymorphism in complexes of ET-binding motifs with viral and host proteins.

**Highlights:**

- The BRD3 ET domain binds to key peptide motifs of diverse host and viral proteins.
- These complexes reveal conformational plasticity in molecular recognition.
- NMR studies demonstrate restricted interdomain motion in the IN CTD / ET complex.
- A cost-effective approach is described for producing isotopically-labeled peptides.

**Etoc Blurb:** We address structurally how the MLV Integrase (IN) usurps the host function of the BET protein through comparative studies of the IN : Brd3 ET complex with that of the host NSD3. MLV integration and thus its pathogenesis is driven through protein interactions of the IN : BET family.

## Introduction

Bromodomain and extra terminal domain containing proteins (BET) scan for epigenetic marks on host chromatin and are also responsible for recruiting large multiprotein complexes to the chromatin. We, amongst others, have shown an important role for host bromodomain and extra terminal domain (BET) proteins BRD2, BRD3 and BRD4 to interact with the MLV IN protein (Aiyer et al., 2014, De Rijck et al., 2013, Sharma et al., 2013, Gupta et al., 2013). The extra terminal (ET) domain of BET proteins interacts specifically with the C-terminal domain (CTD) of MLV IN protein. During MLV infection, the viral integrase associates with the reverse-transcribed viral DNA as part of a nucleoprotein complex called the pre-integration complex (PIC), which also contains other viral proteins including p12 and capsid (CA), along with host proteins including barrier-to-autointegration factor (BAF) (Prizan-Ravid et al., 2010, Fassati and Goff, 1999, Lee and Craigie, 1998). The p12 protein tethers the PIC to mitotic chromatin and ensures nuclear retention through chromatin tethering (Brzezinski et al., 2016b, Elis et al., 2012, Schneider et al., 2013, Wight et al., 2012). However, subsequent interaction with BET proteins is an important determinant for genomic targeting of the PIC towards transcription start sites (TSS) and CpG islands (Wu et al., 2003). More recently it was shown that active promoters / enhancers are the primary integration hotspots for MLV integration (De Ravin et al., 2014, LaFave et al., 2014). Interestingly, the interaction of MLV IN CTD and ET domain can be disrupted by truncation of the terminal 23 amino-acid C-terminal “tail peptide” (TP) region of the CTD (Aiyer et al., 2014). The phenotypic importance of this disruption was evident due to the decreased preference for integration near TSS and CpG islands both *in vitro* (Aiyer et al., 2014, Sharma et al., 2013, De Rijck et al., 2013) and *in vivo* (Loyola et al., 2019).

The ET domain of BET proteins has been described to interact with a multitude of host proteins including NSD3, ATAD5, GLTSCR1, CHD4 and JMJD6 (Rahman et al., 2011, Wai et al., 2018), as well as the Kaposi’s sarcoma herpes virus (KSHV) latency associated nuclear antigen-1 (LANA-1) CTD (Hellert et al., 2013). The mechanism of ET domain directed protein-protein interaction is thought to be conserved across all of these complexes, based on solution NMR studies (Crowe et al., 2016, Zhang et al., 2016, Wai et al., 2018). However, the description of alternative modes of interaction (Konuma et al., 2017) and considerations that previous structural studies have probed only partial interaction interfaces suggest the possibility of plasticity in the molecular recognition mechanisms of ET domains of BET proteins. BET protein inhibitors continue to be assessed for their ability to block a range of cancers (Alqahtani et al., 2019). Accordingly, the ET binding pocket of BET proteins is a novel target site for cancer drug discovery.

Here we describe solution NMR structures of the BRD3 ET domain (residues 554 – 640) in the free form, and in three complexes: (i) bound to the TP (IN_386-407_) of MLV IN CTD, (ii) bound to the complete MLV IN CTD (IN_329-408_), and (iii) bound to a peptide fragment of NSD3 (NSD3_148-184_). This analysis was facilitated by the development of a novel and cost-effective strategy for preparing isotopically-enriched peptides. Combining isotope-enriched TP with isotope-enriched ET provided more extensive NMR data for the TP:ET complex than has been previously available, and a more complete structural characterization. The complex involves formation of an interfacial three-stranded β-sheet, stabilized by interactions with IN residues _400_KIRL_403_, and a hydrophobic interface which includes key IN residue Trp-390. Next, the structure of the larger ∼21 kDa complex of MLV IN CTD with BRD3 ET domain was determined, allowing us to characterize novel structure-function relationships within the CTD of IN, which we classify into three regions: the SH3 domain (residues 329-378), which does not interact with ET, the C-terminal ET-binding motif (ETBM, residues 389-405), which folds into a β-hairpin to form a three-stranded β-sheet with ET, and an intervening partially-flexible linker region (residues 379-388) that tethers IN to ET. ^15^N relaxation studies confirm that the SH3 domain and the ETBM/ET regions of the IN CTD – ET complex are dynamic with respect to each other, but with partially-restricted interdomain flexibility. Finally, the 3D structure of the complex of NSD3_148-184_ with ET was determined, revealing a similar three-stranded β-sheet intermolecular interaction between this ETBM and ET, stabilized by a similar hydrophobic interface and a similar set of complementary electrostatic interactions. However, in this complex the orientation of the β-hairpin in this three-stranded β-sheet is flipped compared to the corresponding IN TP – ET complex. These structures establish a framework for the development of novel inhibitors that can be used for altering retroviral target site selection and disruption of protein-protein interactions, potentially leading to improved cancer therapies.

## Results

Several prior solution NMR structures of complexes formed between ET domains of BET proteins and short peptide motifs of binding partners have described a two-stranded β-sheet interaction (Konuma et al., 2017, Zhang et al., 2016, Wai et al., 2018). In this study, we examine larger viral and host ET binding regions to define their interactions with ET domain, including determinants of hydrophobic and electrostatic interactions. Specifically, we report the NMR solution structure of the BRD3 ET domain alone and in complex with longer polypeptides including the 22-residue MLV IN Tail Peptide (TP), the 79-residue MLV IN CTD, and 37-residue ET-binding region of the host NSD3 protein (Figure 1). These extended polypeptide segments were isotopically enriched using novel fusion protein expression systems. The results and methods described in this study will be broadly applicable in deciphering the role of the ET domain in modulating interaction of different protein complexes with chromatin.

**Fig. 1.**
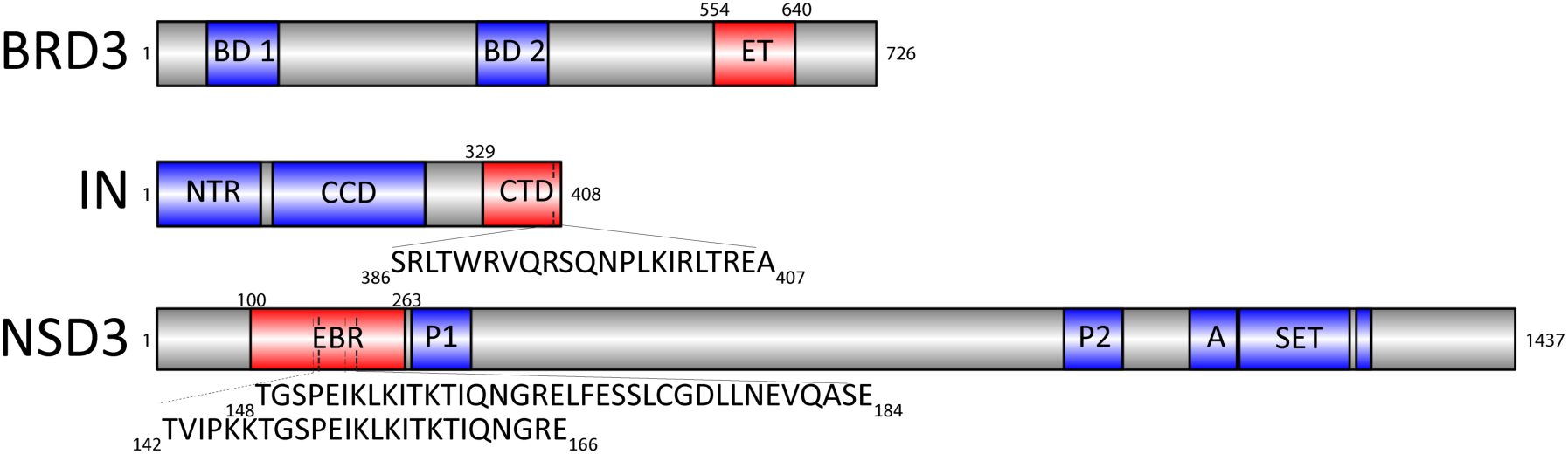
Domain and motif constructs used in this study. The full-length proteins are indicated, with the domains utilized in this study highlighted in red, along with well-annotated domains indicated in blue. BRD3_1-726_ consists of two bromodomains (BD1 and BD2) and the extra terminal (ET) domain. MLV IN_1-408_ consists of the N-terminal region (NTR), catalytic core domain (CCD) and the C-terminal domain (CTD). The sequence of the 22-residue tail peptide (TP) is shown. NSD3 consists of the ET binding region (EBR), two PWWP domains (P), an AWS domain (A), a SET domain, and a post-SET domain. These graphics were created using Illustrator for Biological Sequences (http://ibs.biocuckoo.org/download.php)

### Solution NMR structure of BRD3 ET

Extensive backbone and sidechain resonance assignments were determined for the ∼ 11.2 kDa BRD3 ET_554-640_ domain (Figure 1 and Supplementary S1) using standard triple-resonance NMR methods. An annotated [^15^N-^1^H]-HSQC spectrum is shown in Supplementary Figure S1. The solution NMR structure was determined from 1277 NOE-based conformationally-restricting distance restraints, together with 122 dihedral angle restraints based on backbone chemical shift data, and 38 hydrogen bond restraints identified in intermediate models during the structure analysis process using the program ASDP (Huang et al., 2006) (Supplementary Table S1). The resulting 1437 total experimental restraints correspond to 16.3 restraints per restrained residue. The final structure was refined using CNS in explicit solvent, and the 20 lowest energy structures, of a total of 100 structures calculated, were chosen to represent the solution structure. The resulting structure models have no significant experimental restraint violations. This superimposed ensemble has an atomic root-mean-squared deviation (rmsd) of 0.5 Å for the backbone atoms and 1.0 Å for heavy atoms of well-defined residues. Structure quality assessment scores (Supplementary Table S1) indicate a high quality structure with excellent knowledge-based validation scores (Bhattacharya et al., 2007). The DP score, which assesses the ensemble of models against the NOESY peak list, is 0.78, also indicating a good quality structure (Huang et al., 2012).

The solution NMR structure of BRD3 ET is illustrated in Figure 2, along with its protein sequence and secondary structure. BRD3 ET belongs to the BET pfam domain fold family (El-Gebali et al., 2019), and is conserved amongst other BET proteins including BRD2, BRD3 and BRDT. Rotational correlation time estimates from 1D ^15^N T_1_ and T_2_ measurements (Rossi et al., 2010) indicate that the BRD3 ET is a monomer in solution (τ_c_ = 7.0 ns); this was also confirmed using size-exclusion chromatography (data not shown). Overall this domain is rich in acidic amino acids resulting in a predicted isoelectric point of 4.9. The BRD3 ET structure includes a long unstructured N-terminal region of about 15 amino-acid residues (G554-G569), along with 9 amino-acid residues of the hexahistidine purification tag (Figure 2A). The superimposed ensemble of 20-lowest energy structures is shown in panels B and C as line and ribbon diagrams, respectively. The disordered region at the N and C termini are truncated in the ribbon representation shown in Figure 2C. The BRD3 ET domain consists of three α-helices: α1, residues 574-586; α2, residues 590-602; and α3, residues 624-637 (Figure 2 A-C). The intervening region between α1 and α2 is separated by a kink induced by proline 588, while the α2 and α3 helices are separated by a long loop. This α2 / α3 loop, between helices 2 and 3, is composed of a motif of alternating negatively charged and hydrophobic residues for a stretch of 8 amino acids (_612_DEIEIDFE_619_) (Figure 2D). Surface representation of the hydrophobic residues and electrostatic surface potential reveal the presence of a hydrophobic pocket that is lined on one face by acidic side chains (Figure 2D, E).

**Fig. 2.**
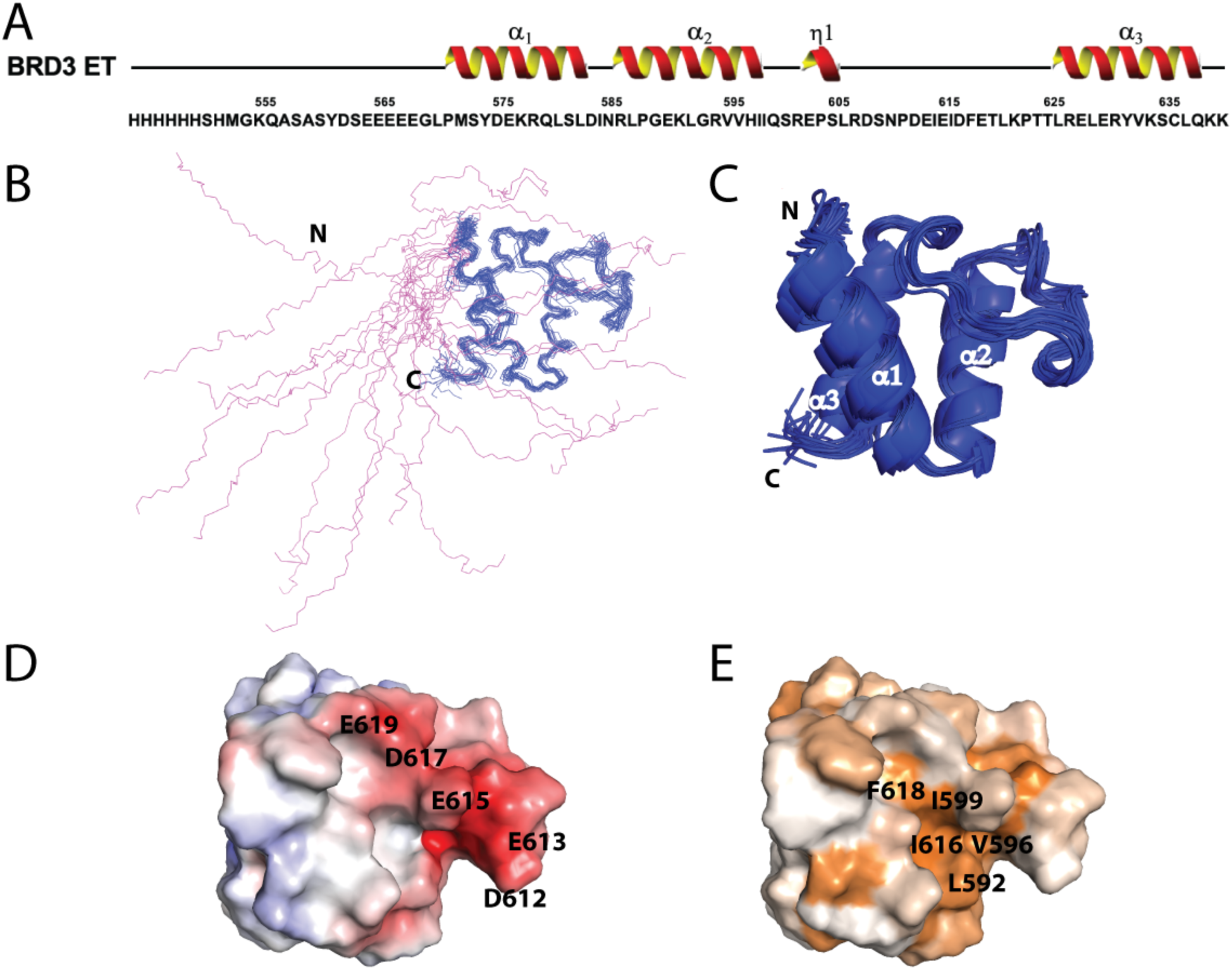
Solution NMR structure of BRD3 ET. (A) Primary sequence of the BRD3 ET construct, including the His_6_ tag. Numbering starts with the BRD3 G554. (B) Representation of the backbone structure of the entire BRD3 ET domain (residues 554-640), including 9 residues of the N-terminal His_6_ tag. The disordered region of the protein is marked in magenta and the ordered region in blue. Further analysis was performed only with respect to the ordered region. (C) Ribbon representation of the ordered region of BRD3 ET domain is shown in blue. The 3 α-helices are α1: 574-586, α2: 590-602, and α3: 624-637. The ordered region of the protein includes residues 569-640. (D) Surface electrostatic representation of the ordered region of BRD3 ET is shown in this panel. Fully saturated red and blue colors represent, respectively, negative and positive potentials of ± 5 kT at an ionic strength of 0.15 M and at a temperature of 298 K. Key negatively charged residues are as indicated. Residues forming the acidic cluster, residues D612-E619, are marked. (E) Surface representation of hydrophobic residues of BRD3 ET reveals the presence of a hydrophobic pocket. Hydrophobicity ranges from brown as most hydrophobic and white as least hydrophobic residues based on hydrophobicity scale (Eisenberg et al., 1984). Key hydrophobic residues are as indicated in the panel.

### Expression and purification of isotopically labeled MLV IN CTD TP

We have previously shown that the MLV IN CTD includes an SH3 fold, together with a long and unstructured TP region comprising of 28 amino acids (Aiyer et al., 2015, Aiyer et al., 2014). Using a 23 amino-acid residue synthetic TP polypeptide, we were able to show the competitive disruption of the BRD3 ET and MLV IN CTD interaction (Aiyer et al., 2014). Production of such small, disordered peptides in *E. coli* generally provides very low yields due to proteolytic degradation in the production process. In order to obtain the solution NMR structure of this IN TP bound to the ET, a novel method of expression and purification of the ^15^N/^13^C-enriched TP was developed (Supplementary Figure S2). We used a C-terminal fusion of an intein-based self-cleavage system to express the peptide as a fusion protein with a non-native methionine at the N-terminus to initiate translation. The peptide consisted of a 22-amino acid segment of MLV IN CTD_386-407_, excluding the terminal Pro-408, since prolines are incompatible with the intein self-cleavage system. Excluding the terminal proline did not affect the stability and binding efficiency of the TP construct (Crowe et al., 2016). On-column self-cleavage of the intein-CBD-6xHis under highly reducing conditions (Mitchell and Lorsch, 2015), resulted in a greater than 90% pure, tagless peptide as assessed by mass spectrometry (Supplementary Figure S2). Using this purification strategy, we obtained highly-purified isotopically-enriched IN CTD TP with a yield of ∼1.5 mgs per 1.5 liter culture. The rapid purification enabled expression of peptide to structure determination within a week, thus providing a highly adaptable platform for solution NMR studies of isotopically-enriched peptides.

In the unbound state, the labeled TP alone showed an [^15^N-^1^H]-HSQC spectrum similar to that of a highly disordered protein (Supplementary Figure S3 A). On addition of unlabeled BRD3 ET domain, there is a marked difference in the [^15^N-^1^H]-HSQC spectrum, indicating disorder-to-order transition (Supplementary Figure S3 B). Owing to the size of the peptide and nature of interaction, resonance assignment of the isotope-enriched peptide bound to unlabeled ET was unambiguous and relatively straightforward, using standard triple resonance experiments. Using isotope-enriched TP with both isotope-enriched and unenriched samples of ET, we then determined the structure of the entire IN CTD TP : ET complex.

### Structure of the MLV IN CTD TP in complex with BRD3 ET domain

The solution NMR structure of the 14.0 kDa complex formed between the MLV IN CTD TP and BRD3 ET was determined from 1533 NOE-based distance restraints, together with 166 dihedral angle restraints, and 22 hydrogen bond restraints (Supplementary Table S1). The resulting 1721 total experimental restraints correspond to 16.2 restraints per restrained residue. The final structure was refined using CNS in explicit solvent, and the 20 lowest energy structures, of a total of 100 structures calculated, were chosen to represent the solution structure. This superimposed ensemble has an atomic rmsd of 0.6 Å for the backbone atoms and 1.1 A for heavy atoms of well-defined residues. Structure quality assessment scores (Supplementary Table S1) indicate a high quality structure with excellent knowledge-based validation scores (Bhattacharya et al., 2007). The DP score is 0.71, also indicating a good quality structure (Huang et al., 2012).

The amino acid sequence, secondary structure, and NMR structure of the BRD3 ET in complex with the MLV IN TP are shown in Figure 3. In this complex, the MLV IN TP adopts an anti-parallel β sheet separated by a turn, which is in part mediated by the conserved Pro398 (Figure 3B and Supplementary Figure S4). Key intermolecular interactions are summarized in Table 1. The hydrophobic cleft of the BRD3 ET accommodates the hydrophobic residues of the MLV IN TP (Figure 3 C,D). Complementary to the BRD3 ET domain, one of the β strands in TP also possesses alternating charged residues _399_LKIRL_403_ that interleave with acidic residues _613_EIEID_617_ of BRD3 ET to stabilize the three-stranded β-sheet interaction (Figure 3, E-G). This interaction is mediated by complementary charged surfaces along with backbone contacts and hydrophobic interactions of the alternate residues consisting of Leu or Ile. Residues Trp390 and Val392 of β6’ also show additional hydrophobic interactions sealing the pocket at the bottom of the binding interface. This interaction with Trp390 could be important due to its conservation across the retroviruses shown in Supplementary Figure S4.

**Fig. 3.**
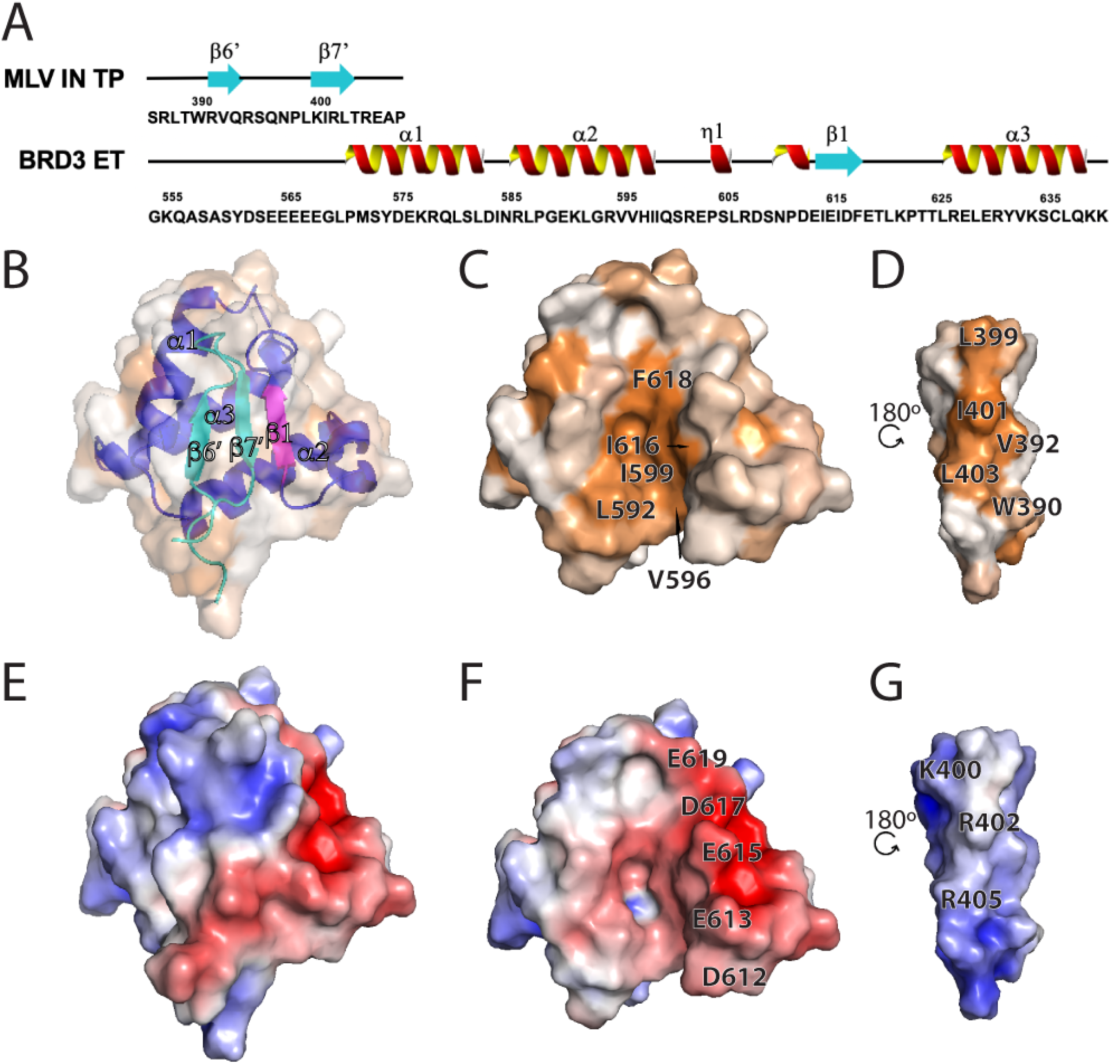
Interaction of MLV IN TP with BRD3 ET. (A) Schematic of the primary sequence of the MLV IN TP and BRD3 ET regions utilized in the studies, together with secondary structural elements as defined in the NMR structures. Here, the N-terminal His_6_ tag of BRD3 ET is not shown. (B) Surface hydrophobic representation of the complex with panel B and panel C oriented in the same plane, and panel D rotated along the y-axis by 180°. Panel B is set at 50% transparency and also shows ribbon representation of the complex with blue/magenta color showing BRD3 ET and cyan color showing the MLV IN TP. The surface hydrophobicity coloring scheme is the same as in Figure 2E. Panel C is the buried face of the BRD3 ET and panel D, the buried face of the MLV IN TP. Key residues forming the hydrophobic pocket are indicated. (E) Surface electrostatic representation of the complex with the E, F and G panels having the identical viewing plane as Figure 3 panels B-D. The surface electrostatic potential coloring scheme is the same as in Figure 2D. Panel F is the surface electrostatic of BRD3 ET and panel G is that of the MLV IN TP. Key residues forming intermolecular electrostatic interactions are indicated.

**Table 1.** Key intermolecular interactions between peptides and BRD3 ET in complexes^a^.

| Intermolecular interactions | BRD3 ET- TP | BRD3 ET- NSD3<sub>148-184</sub> |
| --- | --- | --- |
| Hydrogen bond pairs | Gly589(O), Trp390(NE1);<br>Ile614(O), Leu403(N);<br>Ile614(N), Leu403(O);<br>Glu615(OE2), Ser395(O);<br>Ile616(O), Ile401(N);<br>Ile616(N), Ile401(O);<br>Phe618(N), Leu399(O);<br>Glu619(OE2), Asn397(ND2); | Asp612(O), Lys159(N);<br>Ile614(O), Ile157(N);<br>Ile614(N), Ile157(O);<br>Ile616(N), Leu155(O);<br>Ile616(O), Leu155(N);<br>Phe618(N), Ile153(O);<br>Glu619(N), Ile153(O);<br>Glu619(OE1), Ile153(N); |
| Salt bridges | Asp612(OD1), Arg405(NH2);<br>Glu613(OE1), Arg402(NH1);<br>Glu615(OE2), Arg402(NH2);<br>Asp617(OD2), Lys400(NZ); | Glu613(OE2), Lys156(NZ);<br>Glu615(OE2), Lys154(NZ);<br>Asp617(OD2), Lys154(NZ); |
| Residues of ET forming Hydrophobic interactions | Ile584, Leu592, Val595,<br>Val596, Ile599, Ile614,<br>Ile616, Phe618, Leu621 | Ile584, Leu592, Val595,<br>Val596, Ile599, Ile614,<br>Ile616, Phe618, Leu621 |
| Residues of Peptide forming Hydrophobic interactions | Trp390, Val392, Ile401,<br>Leu403 | Ile153, Leu155, Ile157,<br>Phe168 |

<sup>a</sup>The H-bond and salt bridges were identified using the default settings in CHIMERA (Petterson et. al 2004) and distance measurements using PyMOL with donor-acceptor distances $\leq 3.2$ Å and $\leq 4.0$ Å respectively. Hydrophobic interactions were identified using PyMOL (The PyMOL Molecular Graphics System, Version 2.0 Schrodinger, LLC.).

Interestingly, superposition of [^15^N-^1^H]-HSQC spectra of the bound and unbound BRD3 ET domain shows minor chemical shift perturbations throughout the entire ET domain (Supplementary Figure S5 A, C, D). While the N-terminal disordered segment (residues 554-571) exhibits little or no change upon complex formation, amide ^15^N-^1^H resonances throughout most of the rest of the domain exhibit significant chemical shift perturbations (Δδ N,H > 0.05 ppm, Supplementary Figure S5 C, D). These chemical shift perturbation data indicate global, subtle, allosteric changes throughout the ET structure upon complex formation.

### Interaction of the MLV IN CTD and BRD3 ET domain is restricted to the TP region

In order to determine whether other regions of the CTD SH3 fold are involved in interacting with the ET domain, we next determined the 3D structure of the 21.3 kDa heterodimeric structure of the complex formed between the MLV IN CTD and BRD3 ET. The structure was determined from 1996 NOE-based distance restraints, together with 212 dihedral angle restraints, and 56 hydrogen bond restraints (Supplementary Table S1). The resulting 2264 total experimental restraints correspond to 15.3 restraints per restrained residue. The final structure was refined using CNS in explicit solvent, and the 20 lowest-energy structures, of a total of 100 structures calculated, were chosen to represent the solution structure. In this structure (Figure 4), the SH3 domain and the complex of the ET-binding motif (ETBM) of IN CTD with ET are separated by a 10-residue linker, and the overall structure cannot be superposed due to variability in interdomain orientations. However, the SH3 domain region of IN CTD can be superimposed with rmsd of 0.5 Å for the backbone and 0.8 A for heavy atoms of well-defined residues, and the complex of the IN CTD ETBM with ET can be superimposed with rmsd of 1.1 Å for the backbone atoms and 1.6 A for heavy atoms of well-defined residues. Structure quality assessment scores (Supplementary Table S1) indicate a good quality structure with excellent knowledge-based validation scores (Bhattacharya et al., 2007). The DP score is 0.76, also indicating a good quality structure (Huang et al., 2012).

**Fig. 4.**
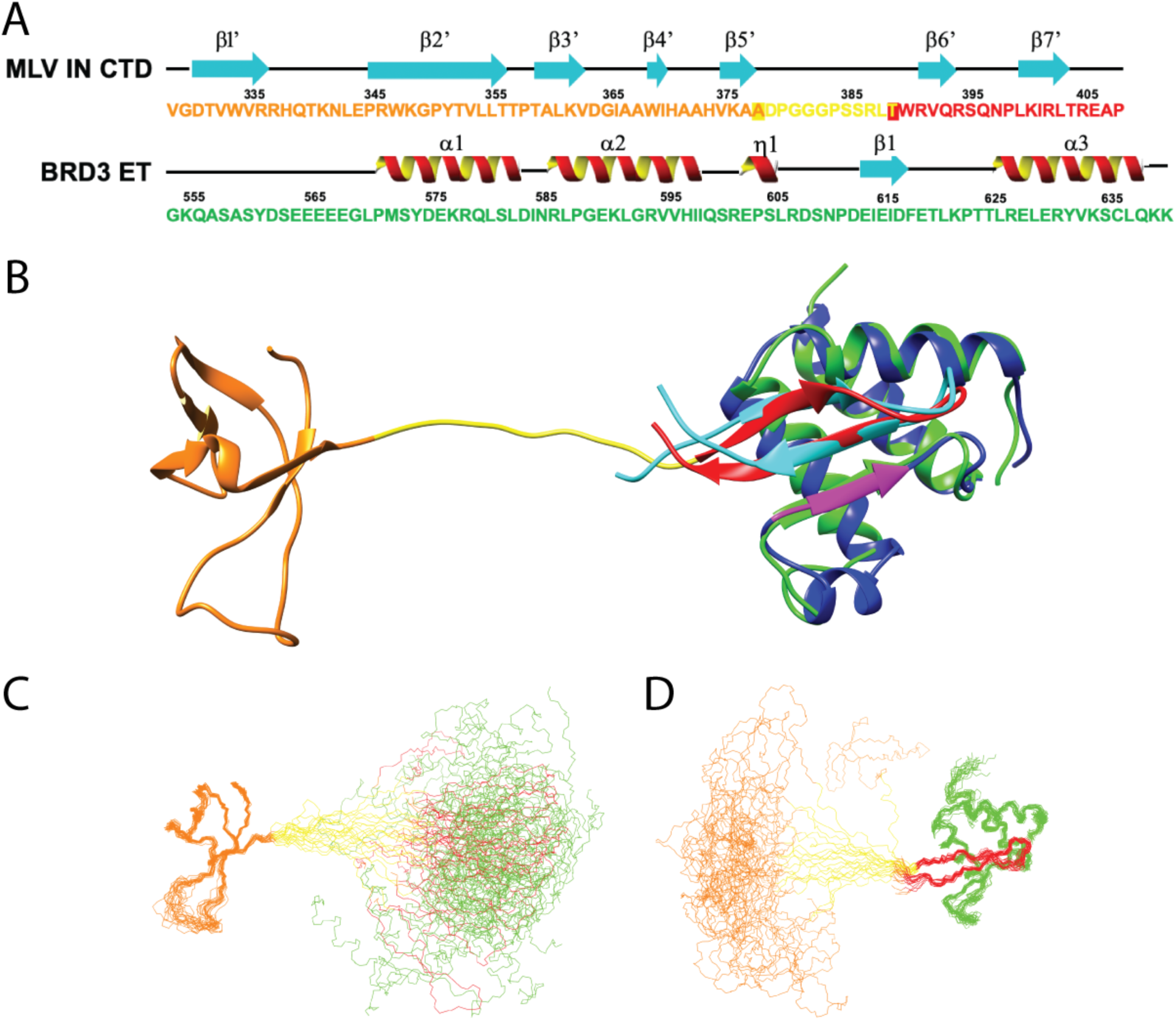
Linker flexibility between SH3 fold and TP of IN CTD it its complex with ET. (A) Schematic of the primary sequence of the MLV IN CTD and BRD3 ET regions utilized in these studies, with the corresponding secondary structural elements as defined in the NMR structures. The SH3 fold is shown in orange, the linker between SH3 fold and TP in yellow, the structured region of the ETBM in red, and the ET domain in green. Transition residues between the SH3 / linker and linker / ETBM are boxed (see Fig. 5D). (B) Ribbon representation of the full CTD-ET complex with color scheme described in A. The ET-peptide structure (blue / magenta / cyan) is overlaid onto the full complex. (C) Backbone representation of the NMR ensemble for the complex, superimposed on the CTD SH3 fold region. (D) Backbone representation of the NMR ensemble for the complex, superimposed on the ETBM : ET domain. In these superimpositions, the SH3 fold is separated from the ETBM by a 10 amino-acid residue linker sequence. Coloring scheme in panels C and D is the same as in panel B. In all three panels the 8 amino acid non-native tag sequence of the CTD and the disordered region of the ET domain (residues 554-568), including the 8 amino acid non-native tag sequence are not shown for clarity.

The NMR structure of the IN CTD : BRD3 ET complex is illustrated in Figure 4, along with the corresponding protein sequences and secondary structures. The overlay of the TP : BRD3 ET structure (ET, blue and magenta; TP, cyan) on the IN CTD : BRD3 ET structure (ET, green; TP, red) shows excellent superimposition of the IN ETBM regions and ET domains of these two structures (Figure 4B). No changes were observed within SH3 fold compared to the NMR structure of the free form of IN CTD in the absence of the ET domain. There are no detectable NOEs between the SH3 domain and the ETBM : ET parts of the complex. Most importantly, the chemical shifts of SH3 in the complex are identical to those of SH3 in free CTD, and the chemical shifts of ET are the same in the TP : ET complex as in the CTD – ET complex (Supplementary Figure S6). These experimental data indicate that the ETBM region of IN alone mediates the interaction between the CTD and BRD3 ET.

In this structure, the orientations of MLV IN SH3 domain and ETBM : ET domain regions of the complex are not well defined with respect to each other. In forming this complex, an approximately 10 residue linker region (D379-L388) is formed between the SH3 fold and the ETBM regions of MLV IN CTD. Flexibility of this linker region can help in facilitating the strand transfer activity of integrase in the nucleus. Disorder prediction analysis (Huang et al., 2014) indicates that this “linker” region has high propensity to be disordered, relative to the rest of the protein (data not shown). In aligning the backbone atoms of the 20 lowest energy conformers of SH3 fold region (Figure 4C), the ETBM : ET domain orientations are highly variable. Conversely, in aligning the ETBM : ET complex, the relative positions of the SH3 fold are highly variable (Figure 4D). The linker region acts as an “tether” between the two domains (Figure 4B,C,D, yellow region). In these structures, the SH3 fold remains independent of the ET domain of the BRD3 protein, separated by the 10 residue linker region.

### Analysis of flexibility / rigidity of the linker region between the IN CTD and TP

Sequence alignment of a panel of gammaretroviruses indicates that the linker region is not highly conserved (Supplemental Figure S4) and varies in size between 6-19 aa between the highly conserved residues Ala378 and Trp390. Additionally, deletion studies of MLV revertants in this region (Loyola et al., 2019) indicated that non-viral sequences can be substituted for 3 of the 7 amino-acid residues maintained in this region (Δ20). To further analyze the dynamics of the linker region, we acquired and analyzed ^15^N nuclear relaxation and ^1^H-^15^N heteronuclear NOE data for the IN CTD : ET complex, which were used to assess variations in internal dynamic motions across the complex and to estimate the rotational correlation times (τ_c_) of the SH3 domain and the ETBM : ET complex of the IN CTD : ET complex. The results of this analysis are shown in Figure 5. The horizontal bars shown in Fig. 5B are the τ_c_ values estimated for these individual domains at 25 °C based on their molecular weights (Rossi et al., 2010); i.e. 4.1 ns for the SH3 domain and 8.4 ns for ETBM : ET complex. These τ_c_ values are consistent with values estimated at 25 °C for the SH3 in the larger IN CTD construct (τ_c_ = 7.0 ns) (Aiyer et al., 2014) and for the ET alone (τ_c_ = 7.0 ns). Remarkably, the overall rotational correlation times τ_c_ for ordered residues of the SH3 domain (orange; τ_c_ ∼ 12.0 ns) and ETBM:ET complex (red + green; τ_c_ ∼ 13.0 ns) components of the IN CTD : ET complex are about the same, and much longer than values measured for these individual domains. In particular, the τ_c_ values for the SH3 domain in the complex (∼ 12 ns) are much higher than expected if the two domains moved independently (4 – 5 ns). Rather, both the SH3 and ETBM:ET regions of the IN CTD : ET complex have rotational correlation times τ_c_ approaching the theoretical value of ∼ 13.3 ns expected for a rigidly tumbling globular complex of 21 kDa at 25 °C (Aiyer et al., 2014). Overall, these results indicate an unexpected strong coupling of the tumbling rates of the two domains in the complex, despite the fact that there are no direct interactions between the domains. These ^15^N T_1_ and T_2_ relaxation rates and [^1^H-^15^N] heteronuclear NOE measurements (Figure 5) also indicate that the linker itself has limited flexibility. While the average τ_c_ value for ordered residues in the IN CTD : ET complex (i.e., excluding the linker region and the flexible N-terminal region of the ET domain) is 12.3 ± 1.5 ns, the linker region has an average τ_c_ of 7.8 ± 1.2 ns. Similarly, the average HetNOE value for ordered residues of CTD-ET complex is 0.74 ± 0.05, while the average HetNOE value in the linker region is 0.26 ± 0.02. These nuclear relaxation measurements are consistent with some degree of stiffness in the “flexible” linker region between the SH3 and ETBM : ET domains of the complex.

**Fig. 5.**
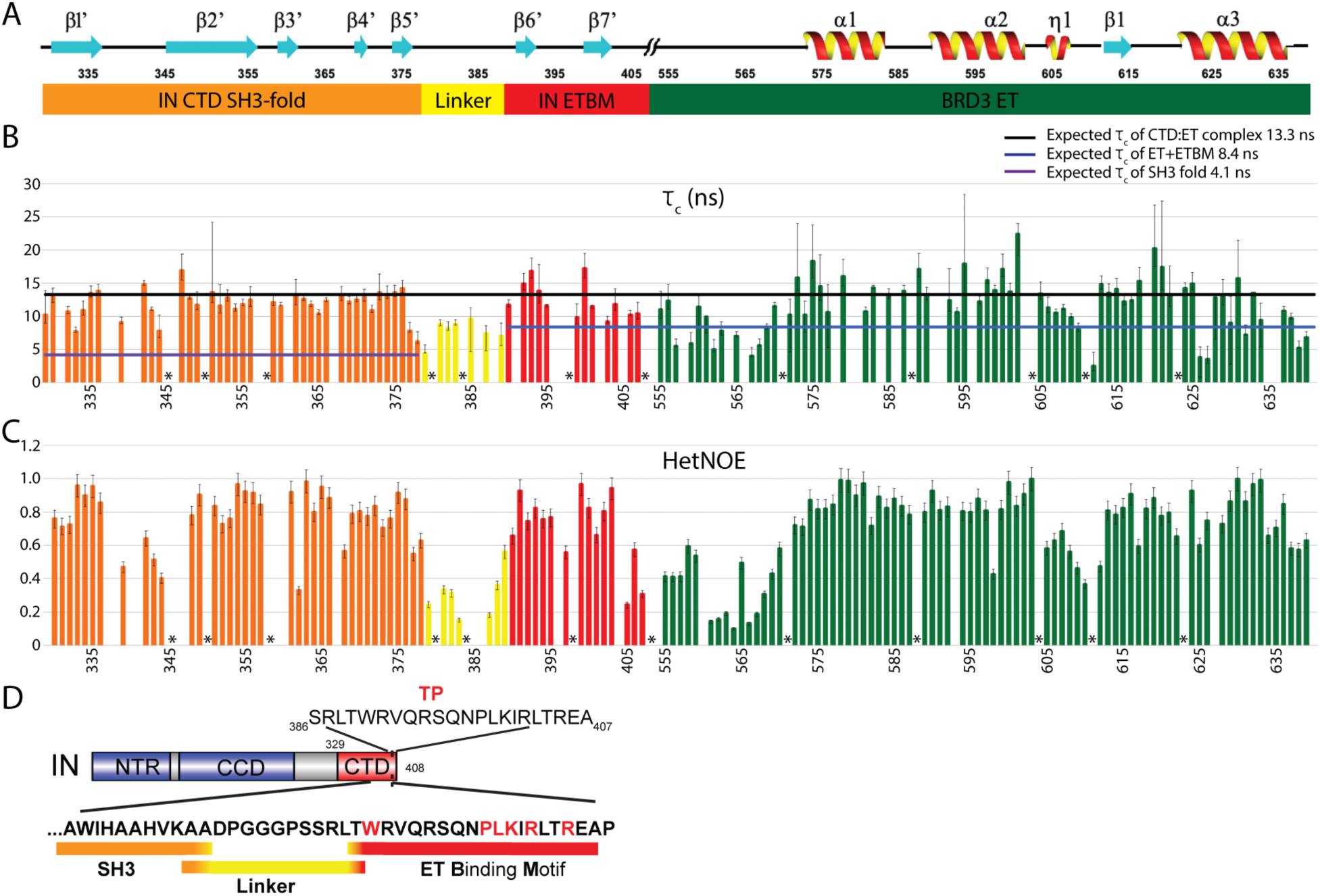
Analysis of chemical shift perturbations and ^15^N-^1^H heteronuclear NOE data for the MLV IN CTD : BRD3 ET complex. (A) Secondary structure features correlating with the amino-acid residue positions in the X-axis in panel B and C. (//) indicates the junction between the C-terminus of the IN CTD and the N-terminus of the BRD3 ET domain. Numbering corresponds to the individual domains in the corresponding full-length protein sequences. Domain/motif coloring scheme: Orange, MLV IN CTD SH3 fold; Yellow, MLV IN CTD linker region; Red, MLV IN CTD ETBM; Green, BRD3 ET domain. (B) N-H bond rotational correlation time measurements (τ_c_ in ns) for each residue across the MLV IN CTD and BRD3 ET complex. Purple horizontal line, overall τ_c_ (4.1 ns) expected for freely-mobile SH3 domain (10.1 kDa) at 25 °C. Blue horizontal line, overall τ_c_ (8.4 ns) expected for freely-mobile ETBM : BRD3 ET complex (14.2 kDa) at 25 °C; Black horizontal line, overall τ_c_ (13.3 ns) expected for rigid MLV IN CTD : ET complex (21 kDa), assuming a rigid body at 25 °C. (C) ^1^H-^15^N Heteronuclear NOE signal (I_saturated_/I_equilibrium_) calculated for each residue across the MLV IN CTD : BRD3 ET complex. Positions of individual amino acids are indicated. (D) Summary of motif boundaries. The sequence of the MLV IN Tail Peptide (TP) used in the NMR structural studies is indicated (top). The SH3 fold (orange) transitions into the linker region at Ala377 and Ala378, which maintain higher HetNOEs compared to their τ_c_ values. The ten amino-acid residues within the Linker region are indicated in yellow. Thr389 transitions the Linker into the ETBM (red). Residues labeled with * correspond to Prolines; no values are plotted for residues which were not assigned, with poor signal-to-noise ratios, and/or with poor relaxation curve fits.

These nuclear relaxation measurements also provide data with which to define the boundaries of the linker region in the IN CTD : ET complex (Figure 5D). The SH3 fold transitions into the linker region starting at residues Ala377/Ala378, with both residues maintaining higher HetNOEs but lower τ_c_ values. Interestingly, in PERV-A, GaLV and KoRV-A integrases, residue Ala377 is replaced with Pro, supporting this transition to linker after Lys376 (Supplementary Figure S4). Similarly, the linker region transitions to the ETBM at residue Thr389, with the ETBM starting with the highly conserved Trp390. These data define the boundaries of the linker region of MLV IN CTD, and identify the transition residues between the SH3, linker and the ETBM (Figure 5D). Overall, these studies identify two functionally-distinct intrinsically-disordered regions of the IN protein. The first intrinsically-disordered region serves as a partially flexible linker that separates the catalytic and assembly functions of the viral protein : DNA integration intasome from the second intrinsically-disordered region that undergoes disorder to order transitions upon binding to the cognate host protein (ETBM).

### Key hydrophobic residue in an NSD3 peptide motif important for stable interaction with BRD3 ET

To extend our interest in defining ET binding domains in the context of full-length or protein subdomains, three constructs of the host NSD3 protein were also generated and analyzed (Figure 1). Initial analysis compared the entire NSD3_100-263_ ETBM with the peptide NSD3_142-166_, which encompasses a previously published minimal ET binding peptide NSD3_152-163_ (Zhang et al., 2016). We observed dramatically different chemical shift perturbations on BRD3 ET upon binding NSD3_142-166_ compared to the complex with NSD3_100-263_ (data not shown), suggesting that there are additional uncharacterized interactions in the longer construct. This prompted us to redefine the boundary of the ET interacting region of NSD3 to include hydrophobic residues Leu167, Phe168, Leu172, Leu176 and Leu177, which are analogous to key hydrophobic residues of IN observed in our solution NMR structure of the IN-CTD : ET complex. The redefined peptide motif NSD3_148-184_ showed a chemical shift perturbation profile on ET (Supplementary Figure S5 B, E, F) similar to that observed for NSD3_100-263_, IN CTD, and IN TP. Significantly, similar chemical shift perturbations throughout ET, attributed to subtle allosteric changes throughout the ET domain structure, are observed upon complex formation with IN CTD, IN TP, and NSD3_148-184_ (Supplementary Figure S5). Hence, similar allosteric changes in ET distant from the peptide binding sites occur in all three of these complexes.

Using the same peptide labelling principle outlined above for IN TP, we designed a SUMO Smt3 fusion with NSD3_148-184_. In this case, the peptide is fused to the C-terminus of Smt3 and can be hydrolyzed with SUMO protease Ulp1 (Supplementary Figure S2). The resulting isotope-enriched peptide was then purified by gel filtration chromatography. As observed for the TP, in the unbound state, the labeled NSD3_148-184_ alone has an HSQC spectrum similar to that of a highly disordered protein (Supplementary Figure S7A). On addition of unlabeled BRD3 ET domain, the [^15^N-^1^H]-HSQC spectrum becomes much better dispersed, indicating disorder-to-order transition (Supplementary Figure S7B). Resonance assignment of the labeled peptide bound to unlabeled ET was unambiguous and relatively straightforward, and using isotope-enriched NSD3_148-184_ with both isotope-enriched and unenriched samples of ET, we then determined the structure of the NSD3_148-184_ : ET complex.

The solution NMR structure of the 15.6 KDa complex of NSD3_148-184_ and BRD3 ET was determined from 1609 NOE-based distance restraints, together with 138 dihedral angle restraints, and 52 hydrogen bond restraints (Supplementary Table S1). The resulting 1799 total experimental restraints correspond to 14.3 restraints per restrained residue. The final structure was refined using CNS in explicit solvent, and the 20 lowest energy structures, of a total of 100 structures calculated, were chosen to represent the solution structure. This superimposed ensemble has rmsd of 0.6 Å for the backbone and 1.1 A for heavy atoms of well-defined residues. Structure quality assessment scores (Supplementary Table S3) indicate a high quality structure with excellent knowledge-based validation scores (Bhattacharya et al., 2007). The DP score is 0.88, indicating a very good quality structure (Huang et al., 2012).

Excitingly, the intermolecular interaction in the NSD3_148-184_ : ET domain complex, made with a longer NSD3 construct than previously reported, now formed a 3-stranded anti-parallel β sheet (Figure 6 A, B), where the smaller NSD3_152-163_ construct forms only a 2-strand β sheet (Zhang et al., 2016). A summary of key intermolecular interactions is presented in Table 1. Mapping of the hydrophobic and electrostatic surface for the NSD3_148-184_ : BRD3 ET complex is shown in Figure 6 B-G. Exposing the buried surface (Figure 6C, D), the hydrophobic cleft of BRD3 ET (residues Leu592, Val596, Ile599, Ile616, and Phe618) is observed to interact with NSD3 residues Ile153, Leu155, Ile157 and Phe168. The electrostatic charge distribution in the BRD3 NSD3_148-184_ : ET : complex is shown in Figure 6E, with the components from the individual domains exposed in Figures 6F, G. The alternate acidic residues of the beta-strand formed in BRD3 ET with residues _613_EIEID_617_ (Figure 6F) interleaving with basic residues _153_IKLKI_157_ to form a three-stranded antiparallel beta strand (Figure 6G).

**Fig. 6.**
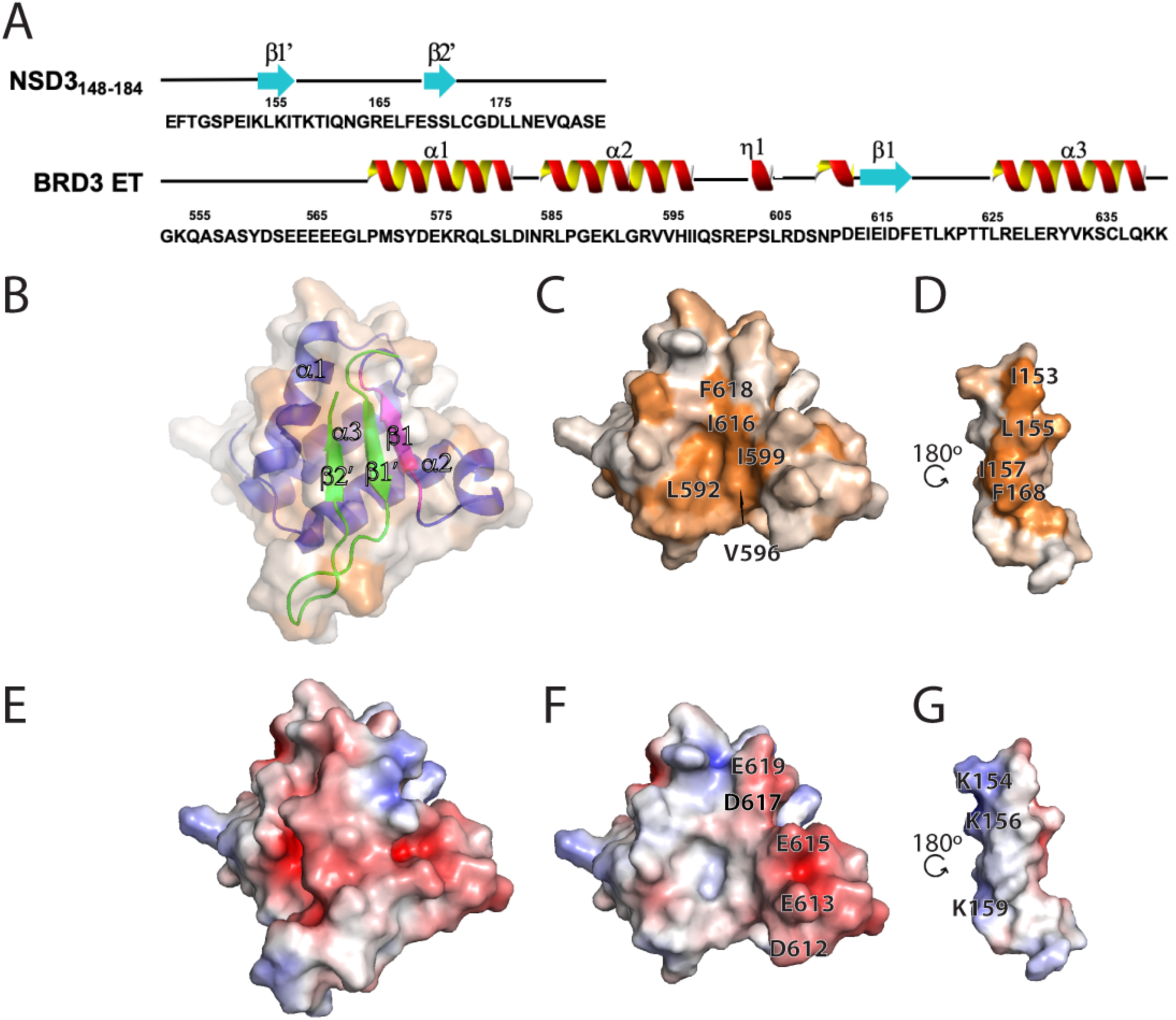
Interaction of BRD3 ET with NSD3_148-184_ peptide. (A) Schematic of the primary sequence of the NSD3**_148-184_** and BRD3 ET regions utilized in the studies, with the corresponding secondary structural elements as defined in the NMR structures. The N-terminal His_6_ tag of BRD3 ET is not shown. (B) Surface hydrophobic representation of the complex with panels B and C oriented in the same plane, and in panel D rotated along the y-axis by 180°. Panel B is set at 50% transparency and also shows ribbon representation of the complex with blue/magenta color showing BRD3 ET and cyan color showing the NSD3**_148-184_**. The surface hydrophobicity coloring scheme is the same as in Figure 2E. Panel C is the buried surface of the BRD3 ET and panel D is that of the NSD3**_148-184_**. Key residues forming the hydrophobic pocket are indicated. (E) Surface electrostatic representation of the complex. Panel F is the electrostatic surface of NSD3 ET and panel G is that of NSD3**_148-184_**. Key residues forming the electrostatic interactions are indicated.

### Alternative binding modes of ET binding motifs

Insights into key biophysical features of the BRD3 ET binding pocket can be obtained through comparisons of the known structures of ET complexes (Figure 7). Figures 7A and B compare the ribbon diagrams of the MLV IN TP and NSD3_148-184_, respectively. Both the NSD3_148-184_ and IN TP : ET complexes involve interactions with the alternating hydrophobic and acidic residues from the ET. However, the register of strands of the NSD3_148-184_ : ET domain complex is different compared to the IN_329-408_ : ET complex (Figure 7C). In the case of the MLV IN CTD, the second β-strand in the C-terminal ETBM interacts with the ET domain, while in the case of NSD3 it is the first strand in the ETBM that interacts with the ET domain (Figure 7C). As a result, the orientation of the peptides forming the β-hairpin in the two complexes are flipped with respect to another. Although the loop between helices α2 and α3 in the ET domain forms a favorable binding pocket for facilitating protein-protein interactions, the mechanism of interaction is distinct for different proteins.

**Fig. 7.**
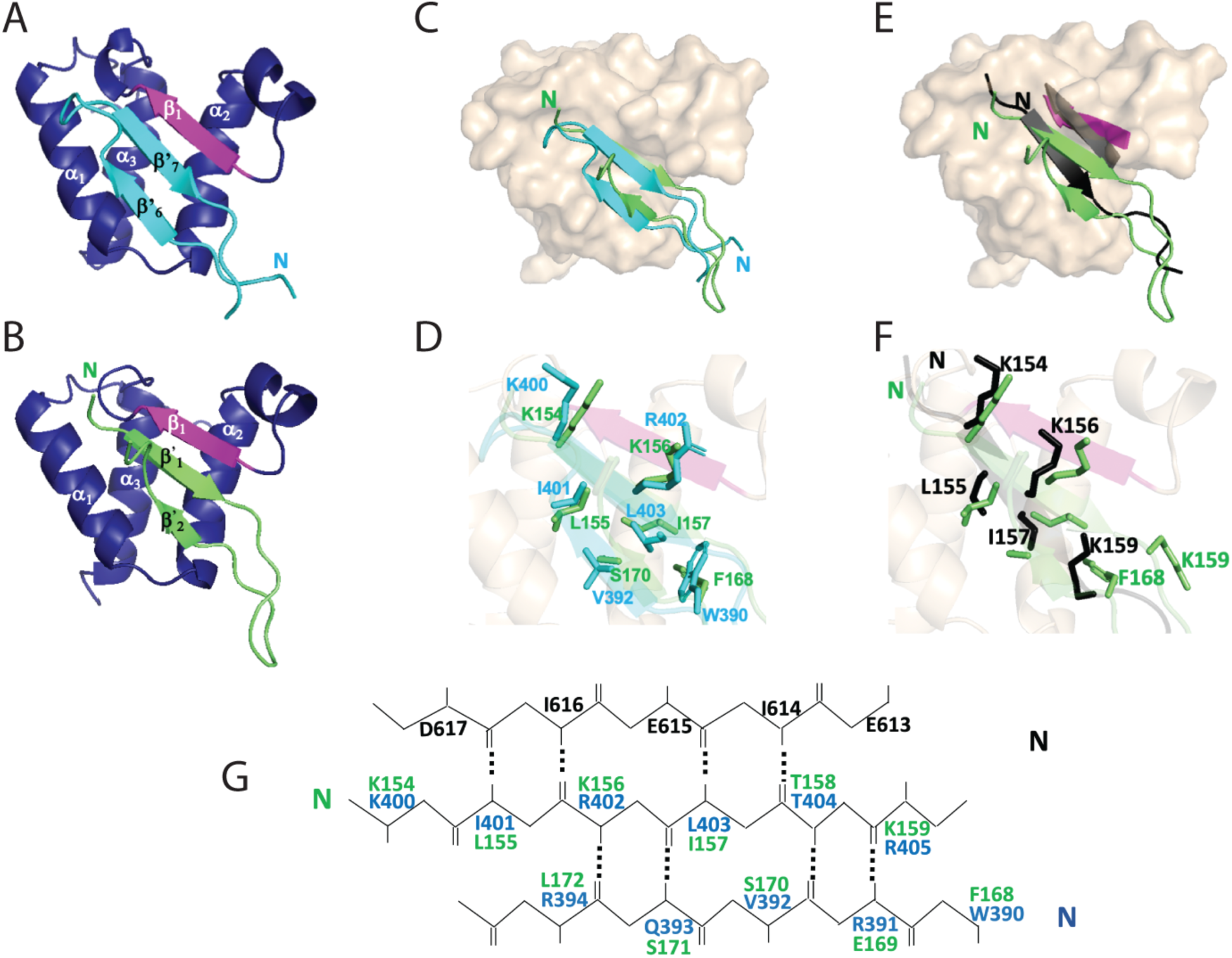
Comparison of peptide : ET complex structures. Coloring of ET domain in complex with the MLV IN TP and NSD3 peptide is as described in Figs 3 and 6. (A) BRD3 ET-MLV IN TP (cyan) (B) BRD3 ET-NSD3_148-184_ (green). (C) Space filling model of BRD3 ET BRD3 ET (beige) overlayed with TP (Cyan) and NSD3_148-184_ (green). (D) Sidechain orientation of key interacting residues of the TP (cyan) with NSD3_148-184_ (green). (E) Comparison of the single beta strand formed using the short NSD3 peptide 152-163 (2ncz) in black and the alignment with the beta strand formed in the ET domain, with the two beta strands formed in the BRD3 ET NSD3_148-184_ complex in green with the alignment of the beta strand of ET in magenta. (F) Sidechain orientations of key interacting residues of the NSD3_152-163_ (black) with NSD3_148-184_ (green) (G) Alignment of key interacting residues of BRD3 ET domain EIEID forming anti-parallel beta strand with TP (residues marked in cyan) and NSD3_148-184_ peptide (residues marked in green).

A superposition of the structures of the complex of BRD3 ET: IN TP and BRD3 ET :NSD3_148-184_, (Figures 7 D and 7 G) indicates that aromatic residue Phe168 of NSD3 occupies the position of Trp390 of MLV IN, sealing the hydrophobic pocket at the bottom. Interestingly, residue Phe168 is conserved in all NSD3 isolates, from Zebrafish to humans (Supplementary Figure S4) as well as between NSD1, NSD2, and NSD3 proteins from humans (Bennett et al., 2017). As can be seen in Figure 7 G, the electrostatic as well as the hydrophobic interactions observed for the TP and NSD3_148-184_ involve analogous residues in the antiparallel beta strand aligned with that formed in the ET domain. The β2’ strand in the latter, however, has the well conserved residue Glu169 that is negatively charged, in contrast to the IN TP which has positively charged residue Arg391 at this position. Interestingly, most gammaretroviruses maintain a positively charged residue at the position; however, both GaLV and KoRV A viruses encode a negatively charged Glu residue at the analogous site, similar to NSD3_148-184_ (Supplementary Figure S4).

Comparison of the extended NSD3_148-184_ : ET domain complex that forms the 3-strand anti-parallel β sheet with the smaller NSD3_152-163_ construct reported to form a 2-strand β sheet. (Zhang et al., 2016) is shown in Figure 7E as a ribbon diagram, with key residues highlight in Figure 7F. For NSD3_148-184_, residues Lys154, Leu155, Lys156, and Ile157 form a first anti-parallel beta stand similar to Zhang et al. (Zhang et al., 2016). However, the beta hairpin is formed with a short strand formed by Phe168, Glu169, Ser170, and Ser171. In the NSD3_152-163_ peptide, Lys159 has the side chain buried into the pocket increasing the hydrophobic interactions of the peptide with BRD4 ET. In the present study with NSD3_148-184_, we find that the aromatic side chain of Phe168 occupies this position and Lys159 provides a secondary interaction with its sidechain. For the BRD3 ET : NSD3_148-184_ complex, the antiparallel beta strand is buried in the hydrophobic pocket while the BRD4 ET : NSD3_152-163_ complex shows the anti-parallel hairpin filling the hydrophobic pocket. The orientation of the ET β1 strand _613_EIEID_617_ is shifted to accommodate the smaller size of the binding NSD3_152-163_ substrate (Figure 7E).

## Discussion

BET proteins such as BRD2, BRD3, BRD4 and BRDT scan for patterns of histone modification on chromatin, through their tandem bromodomains (Taniguchi, 2016, Gilan et al., 2020). Despite structurally similar bromodomains, the tandem bromodomains of BET proteins appear to be functionally distinct (Faivre et al., 2020, Gilan et al., 2020). The ET domain on the other hand predominantly functions in protein-protein interaction networks for recruitment of gene-expression regulating proteins and protein complexes (Rahman et al., 2011, Wai et al., 2018, Konuma et al., 2017, Zhang et al., 2016). The interplay between the functions of the tandem bromodomains and ET domain is likely to give phenotypic distinction to the respective BET proteins. In this study, we report solution NMR structures of BRD3 ET alone and of complexes formed between the BRD3 ET domain with the MLV IN TP_386-407_, the MLV IN CTD_329-408_, and the NSD3_148-184_, allowing for structural comparison of the mechanism of binding of different substrates.

Biochemical studies of MLV virus bearing deletion of the IN TP resulted in loss of BET protein interaction and decreased integration at promoter/enhancer regions (Aiyer et al., 2014, Loyola et al., 2019, Sharma et al., 2013, De Rijck et al., 2013). However, it was noted that integrations at TSS and CpG islands did not reach baseline, implying that alternative secondary ET binding sites within IN remained a possibility (Aiyer et al., 2015). Positions within the MLV IN CCD were proposed to interact with BET proteins based on amino acid substitution studies (Gupta et al., 2013). However, mutagenesis of these residues did not yield viable virus (Loyola et al., 2019) and homology models of the MLV CCD (Aiyer et al., 2015), indicated that the residues important for BET protein interaction are buried and close to the dimer interface of the outer and inner CCDs, thus making it an unlikely interaction interface (Loyola et al., 2019). Within the IN CTD, a mild decrease in preference for TSS and CpG islands was observed for constructs bearing substitutions in the MLV IN CTD β1-β2 loop (Aiyer et al., 2015). This prompted the NMR solution structure analysis of the complete IN CTD_329-408_ in complex with the BRD3 ET domain. Surprisingly, no secondary interaction sites are observed between the ET domain and CTD beyond the IN TP_386-407_ ET-binding motif. This does not rule out the possibility of other regions of the BET protein being involved in secondary interactions, either directly with MLV IN or indirectly through the PIC.

The intrinsically-disordered C-terminal region of MLV-IN has two distinct structure-dynamic function aspects; (i) the final ∼ 20-residues are the ET-binding motif, which undergoes a disorder-to-order structure as part of its intermolecular recognition mechanism, resulting in (ii) a polypeptide linker between this tethered site and the rest of the IN molecule. Our NMR studies further demonstrate that the resulting polypeptide linker of IN has restricted flexibility. Molecular modeling indicates that these dynamics limit the range of orientations of IN with respect to BRD3 in the IN-nucleosome complex.

Linker regions have been increasingly implicated in diverse roles such as modulating propagation of allostery and restricting the sampling space of relevant conformations (Ma et al., 2011, Papaleo et al., 2016, Piai et al., 2016). The absence of contact between the SH3-fold and the rest of the IN CTD-ET complex underscores a function for the linker region separating the two regions. We used our understanding of the structure of the ETBM together with disorder prediction methods (Huang et al., 2014) to analyze the interdomain linker regions that could form in other ET complexes. The size of this region varies among related gammaretroviruses between five residues in PERVs to eighteen in FeLV (Supplementary Figure S4). For M-MLV, the linker region is ten amino-acid residues (_379_DPGGGPSSRL_388_), although deletion mutants have been isolated in mice containing a spacer of six residues, three of which are non-viral in nature but maintaining one of the two Pro residues (Supplementary Figure S4) (Loyola et al., 2019). Within all of the gammaretroviral linker regions, Pro residues are highly abundant. For example, Pro residues account for three of the five residues in the predicted linker sequence of PERV. Additionally, Gly and Ser residues are also common in these linker regions. Variations in composition of Gly and Ser residue in an otherwise intrinsically-disordered linker can affect the stiffness / flexibility of the linker (van Rosmalen et al., 2017). This, plus the unequal linker lengths, make it difficult to predict the degree of flexibility of various gammaretroviral linker regions. Interestingly, the NSD3 ET binding motif is flanked by low-complexity regions [DisMeta, (Huang et al., 2014)], including a Pro-rich sequence N-terminal to the ETBM (Supplementary Figure S4), that are predicted to be intrinsically-disordered.

In order to evaluate the nature of linker flexibility, we used ^15^N nuclear relaxation and ^1^H-^15^N Heteronuclear NOE measurements. The linker region in the MLV IN CTD bound to ET has some flexibility. The conclusion is based on four key observations. First, there are no contacts indicated by NOE or chemical shift perturbation measurements between the SH3 and ET domains of the complex. Second, structure determination results in models in which the relative orientation between the domains is not uniquely determined. Although the NMR methods used to determine the structure of the CTD-ET complex are not suited for determining the dynamic distributions of structures, the NMR data simply do not provide any restraints between the two domains. Third, estimates of the rotational correlation times τ_c_ indicate that in the complex both the SH3 and the ET domains have τ_c_ values that are significantly greater than those of these individual domains, and approaching τ_c_ values expected for a rigid complex. Finally, the HetNOE and τ_c_ values estimated within the linker indicate more order than expected for a highly disordered linker. Collectively, these results indicate that while long flexible linkers are not a prerequisite for successful interaction of IN with BET proteins, the appropriate degree of flexibility in these tethers created by complex formation might play an important role in entropically stabilizing binding of IN with BET protein, thus facilitating the functional targeting of integration sites during gammaretroviral infection.

Recently it was proposed that BET proteins interact with MLV IN after p12-mediated nuclear retention of the viral PIC (Brzezinski et al., 2016a, Brzezinski et al., 2016b, Borrenberghs et al., 2019). Using FRET-based studies, the interaction of BET proteins with MLV IN was shown to cause alteration in the quaternary structure of IN within the PIC (Borrenberghs et al., 2019). An overlay of the IN CTD : BRD3 ET domain onto the PFV intasome provides a hypothetical model of the architecture of the intasome complex with host nucleosomes through BET proteins (Figure 8). The linker region of the docked model of IN CTD and ET domain required alteration of the relative domain orientations in order prevent steric clashes with the rest of the intasome. This suggests that the linker separating integrase and BRD3 interaction can modulate conformational changes within the PIC to engage host chromatin. Since the targeting of intasomes *in vivo* can be altered by truncating the ET binding motif, substituting the ET binding motif to retarget integration may require a linker region that optimally orients the SH3 domain and the retargeting module.

**Fig. 8.**
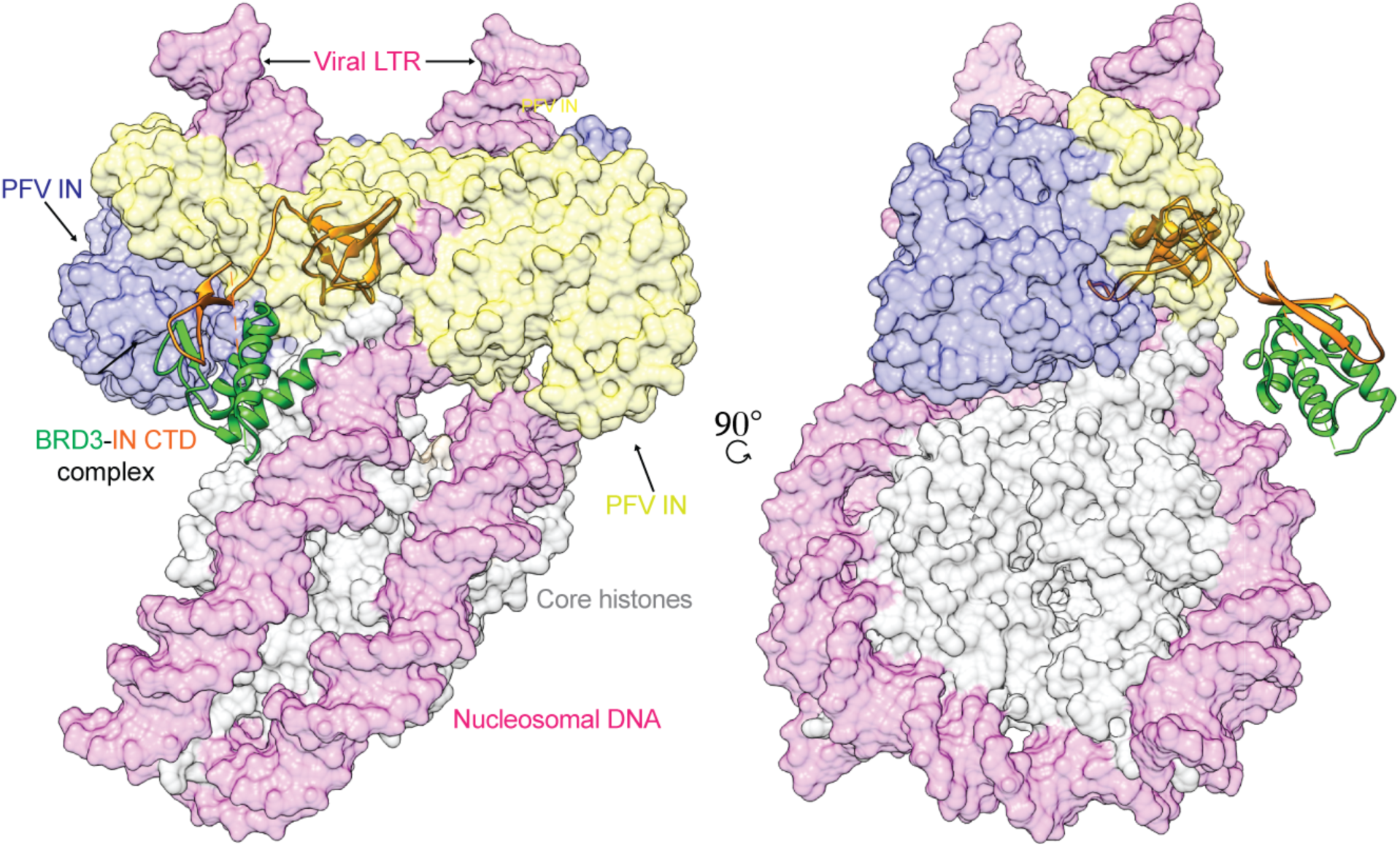
Intasome overlay. Left panel represents the overlay of PFV intasome with the MLV IN CTD and BRD3 ET complex (this study). The alignment of PFV intasome-IN CTD:ET complex onto nucleosome was guided using the PFV intasome-nucleosome cryo-EM structure (PDB ID 6RNY; EMDB ID 4960) (Wilson et al., 2019). The PFV intasome dimer of dimers is represented in yellow and blue color, the IN CTD is in orange, BRD3 ET complex is in green, the viral and target DNA is in magenta and core histone proteins are in grey. Right panel is a 90° rotation along the y-axis.

Maintaining an unstructured region within a protein may be a general feature to facilitate BET protein interaction. In the absence of ET, the MLV IN C-terminal amino-acid residues encoding the linker plus ETBM are unstructured (Aiyer et al., 2015). The CHD4 ET binding domain is localized to its N-terminal region (aa 290-301) and is reported to be unstructured in the absence of ET (Wai et al., 2018). Analysis of the NSD3 binding domain using DisMeta (Huang et al., 2014) indicates this region is also intrinsically disordered, with the propensity to form a beta-sheet in the region of aa 153-160. For the BRD3 ET domain, DisMeta predicts the region surrounding and including the β-strand to be less ordered than the domain as a whole and correlates with the solution NMR structures reported herein of the domain in the absence and presence of either the IN TP or NSD3_148-184_ peptides. For Influenza virus, the flexible linker within its NonStructural protein 1 (NS1) is essential for its pleiotropic functions to bind multiple cellular factors during viral replication (Mitra et al., 2019, Das et al., 2008). This conformational polymorphism is similarly reflected in the ability of the ET-binding cleft to transition disordered binding partners into ordered structures in an orientation independent fashion.

Although there is a strong degree of homology between the ET domains of BRD2, BRD3 and BRD4, there are some reported differences between the different members of BET proteins. Of note, the structure of the BRD3 ET:TP complex is highly homologous to that reported for BRD4 ET: TP (Crowe et al., 2016) and this domain would not provide selective recognition and/or binding between BET family members for MLV IN. An isoform of BRD3 that lacks the entire ET domain called BRD3R is important for nuclear reprograming and mitosis (Shao et al., 2016). To our knowledge, none of the other human BET proteins has isoforms that lack the entire ET domain. Pull-down assays using BRD3 ET as bait did not show interaction with full length JMJD6, but BRD2 and BRD4 on the other hand are capable of interacting with JMJD6 (Rahman et al., 2011). In the case of BRD4 ET, the interacting interface was shown to be restricted to an α-helical region from residues 84-96 in JMJD6 using a peptide motif spanning that region. However, in the full length JMJD6 protein, this region is not completely solvent exposed and was proposed to require allosteric modulation by ssRNA to stabilize the ET:JMJD6 interaction (Konuma et al., 2017). A separate report showed no interaction between BRD4_471-730_, which includes the extended ET domain and JMJD6_1-140_, which includes the purported ET interacting region, indicating the peptide motif (residues 84-96) alone may not be mediating the interaction. In the same report, JMJD6_1-305_ was shown to be minimally required for interaction with BRD4_471-730_ (Liu et al., 2013). We performed [^15^N-^1^H]-HSQC experiments with isotopically enriched BRD3 ET_554-640_ and unlabeled JMJD6_1-336_ (Supplemental Figure S8) and small but detectable chemical shift perturbations similar to those observed for the other complexes reported in this study. This shows the variations between the different BET family members as well as that observed for the full-length JMJD6 proteins versus that observed for a minimal alpha helical peptide.

Our solution NMR structural studies were greatly facilitated through the development of methods to isotopically enrich, express and purify labeled peptides through a cost-effective production scheme. Typically, isotopically labeled synthetic peptides are expensive with 1 mg of labelled peptide costing anywhere from $5000-$10000. Two parallel expression and purification systems were utilized using either a chitin binding domain and intein/DTT cleavage or a SUMO (Smt3) fusion/SUMO protease combination. This approach greatly facilitated solving the structure of the IN TP and NSD3_148-184_ respectively. Solving structures using unlabeled peptides was time consuming as a consequence of repetitive amino acid residues in the sequence (5 Arg in TP and 5 Leu in NSD3_148-184_), making unambiguous resonance assignments challenging. Additionally, many commercial peptide synthesis companies do not provide isotopically enriched tryptophan residues. Since Trp390 in the IN CTD TP was an important residue for stabilizing the interaction, this peptide purification scheme was greatly beneficial in the elucidation of the solution NMR structures of the IN-ET complex. The technique is simple to implement and can be extended to smaller peptides with a reverse-phase HPLC or peptide FPLC size exclusion chromatography for purification and solvent exchange.

Solution NMR structures of complexes involving BRD3 ET and BRD4 ET have been described before with peptides such as ET binding motif, LANA-1 CTD_1133-1144_, NSD3_152-163_, JMJD6_84-96_, BRG1_1591-1602_ and CHD4_290-301_ (Konuma et al., 2017, Zhang et al., 2016, Wai et al., 2018). A key component of the interaction between the BRD3 ET domain and MLV IN CTD TP is the formation of a hydrophobic pocket that stabilizes this interaction. Substitution of a single hydrophobic amino acid in this pocket is capable of disrupting this interaction in the case of MLV IN (De Rijck et al., 2013). Unlike MLV IN TP, prior published structures of the peptide motifs of NSD3_152-163_, LANA CTD_1133-1144_, BRG1_1591-1602_ and CHD4_290-301_ observe only a double-stranded interface between the peptide and the ET domain (Konuma et al., 2017, Wai et al., 2018, Zhang et al., 2016). The extended NSD3_148-184_ peptide forms a three-stranded interaction, which accommodates the binding of residue Phe_168_ into the hydrophobic pocket occupied by Trp_390_ of MLV IN (Figure 7 D, G). Interestingly, the CHD4_290-301_ peptide bound to the ET encodes a terminal Phe in the vicinity of this hydrophobic pocket (Wai et al., 2018). For NSD3, Lys_159_ is conserved across all species (Supplementary Figure S4) and might assist in the initial binding of the region to ET, which is subsequently displaced when Phe_168_ can stabilize the complex. For MLV IN, Arg405 is similarly conserved, however no binding to GST-BRD4 ET of MLV IN_W390A_ was observed using an AlphaScreen (De Rijck et al., 2013). Thus, these studies extend the commonality of the minimal ligand binding by ET beyond the electrostatic recognition (Zhang et al., 2016) of the ET EIEID sequence to now include the secondary interactions within the hydrophobic pocket.

Although the hydrophobic pocket and the acidic residues that line this pocket is a common feature of these protein-protein interactions from the ET side, in this study we have shown that the interacting partners can show diverse structural features. The MLV IN CTD TP and NSD3_148-184_ ET binding peptide both form β-hairpins but in an inverted orientation to one another. Significantly, for both MLV IN and NSD3 the sequences in the turn region (_398_PLK_400_ for MLV and _162_QNGRE_166_ for NSD3, Supplemental Figure S4), are also highly conserved. In all of these structures, the ET domain forms a β-strand involving the loop polypeptide segment between helices α2 and α3. However, the binding motif to ET studied within JMJD6 is reported to be an α-helix (Konuma et al., 2017), showing the potential diversity of structures capable of fitting into this cavity. Of all the available ET structures with bound peptide motifs, only the MLV IN TP complex has a binding constant in the sub-micromolar range (Crowe et al., 2016, Konuma et al., 2017) indicating the extended hydrophobic interactions are important for stable binding. Table 2 analyzes the solvent accessibility using GETAREA/PyMol (Fraczkiewicz and Braun, 1998) of the BRD3 ET in complex with IN TP and NSD3_148-184_ from this study with the BRD4: JMJD6 (PDB ID 6BNH) and NSD3_152-163_ (PDB 2NCZ). This analysis demonstrates that the 3-stranded beta-sheet complexes result in an increased buried surface area, which should result in tighter binding for these extended structures (Konuma et al., 2017, Zhang et al., 2016).

**Table 2.**
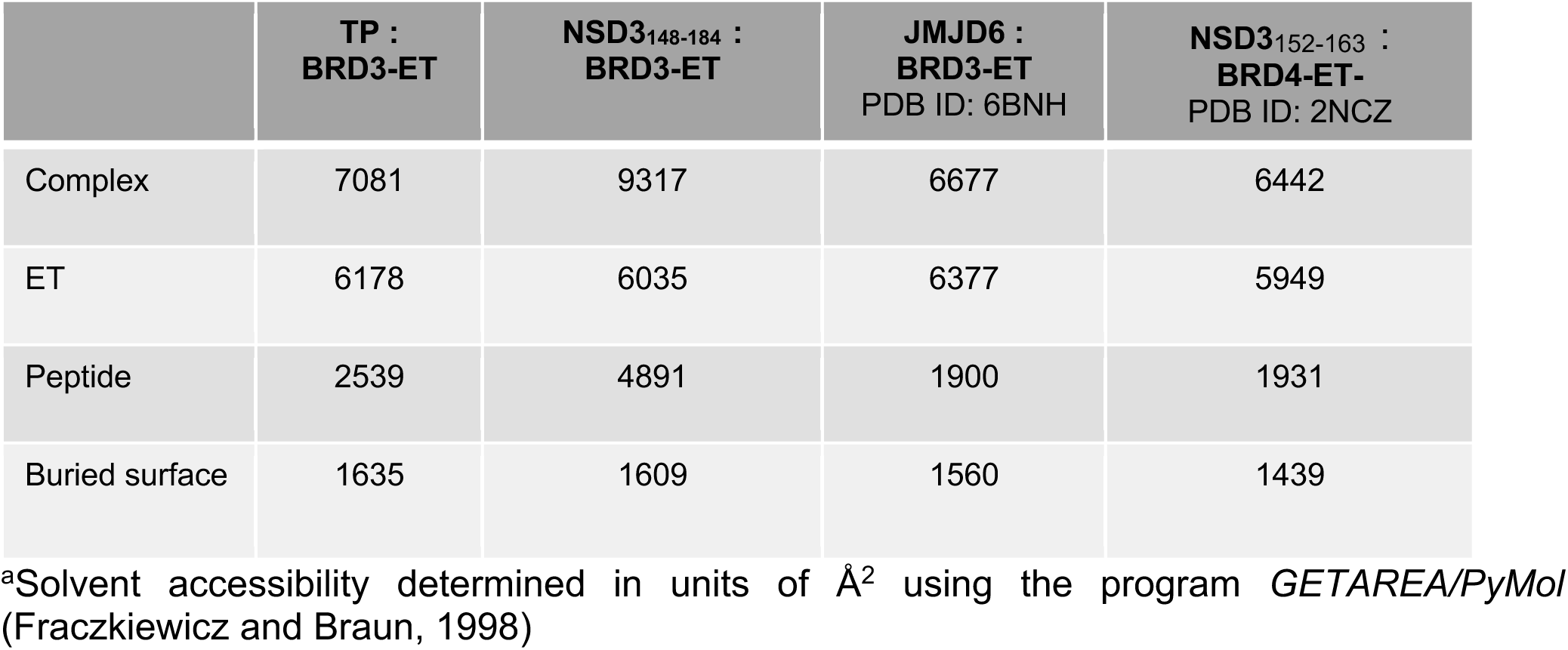
Buried surface analysis for interfaces of BRD ET complexes^a^.

A general caveat in using really short peptide motifs that were designed based on primary sequence alone as in LANA, NSD3, CHD4, BRG1 and JMJD6 (Wai et al., 2018, Konuma et al., 2017, Zhang et al., 2016) is that those interactions need not necessarily translate in the context of full-length complexes. In the case of LANA-1 CTD, the peptide motif from 1133-1144 was shown to be sufficient to interact with BRD4 ET domain with the three C-terminal residues not involved in extensive interactions (Zhang et al., 2016). Studies involving larger complexes of the entire LANA CTD and BRD2/4 ET domains showed distinct regions being important for the stability of these complexes with the 1125-1129 region in LANA CTD showing the highest chemical shift perturbation upon ET binding (Hellert et al., 2013). We designed an extended peptide encompassing residues 1108-1141 that combined features of both the Zhang et al and Hellert et al studies (Hellert et al., 2013, Zhang et al., 2016). However, we observed only a weak interaction with BRD3 ET and LANA-1 CTD_1108-1141_ (Supplementary Figure S7), suggesting additional interfaces like the ‘basic top’ are needed between the two domains and possibly the oligomeric nature of LANA-1 CTD (Hellert et al., 2013) can modulate stable interactions.

Currently, BET inhibitors used as potential cancer therapeutics and/or tool reagents target only the bromodomains of the BET family of proteins, either individually or in tandem (Alqahtani et al., 2019). Despite their efficacy and being clinically promising targets, bromodomain inhibitors are associated with development of drug resistance mechanisms and other toxicities (Alqahtani et al., 2019, Jin et al., 2018). There is an increasing need to tune therapeutic strategies targeting epigenetic readers like BET proteins, since their activity is highly tissue specific. The ET domain provides a flexible interacting interface enabling interaction with various host and viral proteins. This interacting interface thus provides a highly context dependent mode for the different interacting partners and can serve as a plausible drug target. NSD3 is frequently amplified or occurs as fusions in a variety of human cancers and predominantly perturbs cell cycle progression pathways (Han et al., 2018). Many of these cancers involve the BRD4-MYC or BRD4-CHD8 axis regulating super-enhancer activity of oncogenic transcriptional network (Loven et al., 2013, Han et al., 2018). Since BRD3 and BRD4 both interact with NSD3, the extended BRD3 ET : NSD3_148-184_ interface can be used as a starting platform, to design therapeutic strategies to disrupt the interaction. We believe that the plasticity of interaction mechanisms of the ET domain with other proteins can extend this strategy to serve as a platform for developing targeted therapeutic responses towards certain forms of cancers with BET protein vulnerabilities.

## Acknowledgment

This work is supported by grants from NIH grants R35 GM122518, R01 GM110639 (MR) and R01 GM120574 (GTM).

## Author Contributions

Conceptualization, MR, GTM, SA, GVTS; Methodology, GTM, GVTS, GL, GC and SA; Software, GVTS and GC; Validation, SA, GVTS, GL, JH, GC, and GTM; Formal Analysis, SA, GVTS, JH, and GTM,; Investigation, SA, LM, BCJ, and GVTS; Resources, SA, LM, and GTM; Data curation, GVTS and SA; Writing-original draft, SA, GVTS, JH, GTM and MR; Writing, Review and Editing: SA, GVTS, LM, JH, BCJ, GTM, and MJR; Visualization, SA, GVTS, GL, JH, GTM and MJR; Supervision, GTM and MJR; Project Administration, GTM and MJR; Funding Acquisition, GTM and MJR.

## Star Methods

### Key Resources

Star Methods Table SM1.

**Star Methods**1053 **Key Resources – Star Methods Table SM1**
| REAGENT or RESOURCE | SOURCE | IDENTIFIER |
| --- | --- | --- |
| **Bacterial and Virus Strains** |  |  |
| *E. coli* BL21(DE3) | Novagen | Roth lab |
| **Chemicals, Peptides, and Recombinant Proteins** |  |  |
| MLV IN CTD<sub>386-407</sub> peptide | Peptide 2.0 Inc | N/A |
| NSD3<sub>148-184</sub> peptide | Peptide 2.0 Inc | N/A |
| LANA<sub>1108-1141</sub> peptide | Peptide 2.0 Inc | N/A |
| Sodium chloride | Sigma Aldrich | 746398-2 |
| Tris | Sigma Aldrich | T1503 |
| TCEP | GoldBio | #TCEP1 |
| Sodium phosphate | JT Baker/Fisher | 3828-05/S381-3 |
| 2-mercaptoethanol | BioRad | 161-0710 |
| Imidazole | JT Baker | N811-06 |
| <sup>15</sup>N-labeled ammonium chloride | Cambridge Isotope | #NLM-713-PK |
| <sup>13</sup>C-glucose | Cambridge Isotope | #CLM-1396-PK |
| NdeI | NEB | R0111L |
| SpeI | NEB | R0133S |
| EcoRI HF | NEB | R3101T |
| HindIII HF | NEB | R3104T |
| EDTA | JT Baker | 8993-01 |
| Dithiothreitol | Fluka Analytical | 43819-5G |
| Ni-NTA resin | Qiagen | 30430 |
| **Biological Samples** |  |  |
| 143BCeB cDNA | 143B provided by F. González-Scarano, U. Pennsylvania, Philadelphia, PA) (Bupp et al., 2005) | Roth lab |
| **Recombinant DNA** |  |  |
| pCeB | Provided by Y. Takeuchi. (Cosset *et al.*, 1995b) |  |
| pGV58 MLV IN CTD<sub>386-407</sub> | This paper | Roth and Montelione labs |
| pET15NESG MLV IN CTD<sub>329-408</sub> | Aiyer et al | Roth and Montelione labs |
| pET15NESG BRD3 ET | Aiyer et al | Roth and Montelione labs |
| pSUMO NSD3<sub>148-184</sub> | This paper | Roth and Montelione labs |
| pGV3358 LANA<sub>1108-1141</sub> | This paper | Roth and Montelione labs |
| pSUMO JMJD6<sub>1-336</sub> | This paper | Roth and Montelione labs |
| **Deposited Data** |  |  |
| BRD3 ET domain. Solution NMR structure coordinates, NMR resonance assignments, extensive raw FID data and NOESY peak lists. | This paper | PDB: 7JMY; BMRB: 30782 |
| BRD3 ET domain in complex with MLV IN CTD TP. Solution NMR structure coordinates, NMR resonance assignments, extensive raw FID data and NOESY peak lists. | This paper | PDB: 7JQ8; BMRB: 30786 |
| BRD3 ET domain in complex with MLV IN CTD. Solution NMR structure coordinates, NMR resonance assignments, extensive raw FID data and NOESY peak lists. | This paper | PDB: 7JYZ; BMRB: 30791 |
| BRD3 ET domain in complex with NSD3<sub>148-184</sub>. Solution NMR structure coordinates, NMR resonance assignments, extensive raw FID data and NOESY peak lists. | This paper | PDB: 7JYN; BMRB: 30790 |
| **Oligonucleotides** |  |  |
| Primers for cloning MLV IN CTD<sub>386-407</sub> in pGV358, see Table S1 | IDT |  |
| Gene block for NSD3<sub>100-263</sub> in pSUMO, see Table S1 | IDT |  |
| Primers for cloning NSD3<sub>148-184</sub> in pSUMO, see Table S1 | IDT |  |
| Primers for cloning LANA<sub>1108-1141</sub> in pGV358, see Table S1 | IDT |  |
| Primers for cloning JMJD6<sub>1-336</sub> in pSUMO, see Table S1 | IDT |  |
| Primers for cloning NSD3<sub>142-166</sub> in pSUMO, see Table S1 | IDT |  |
| **Software and Algorithms** |  |  |
| Topspin | Bruker Biospin | www.bruker.com |
| CYANA | (Guntert et al., 1997, Herrmann et al., 2002) |  |
| Sparky | (Lee et al., 2015) |  |
| PyMOL | PyMOL Molecular Graphics System | pymol.org |
| UCSF Chimera | (Pettersen et al., 2004) |  |
| NMRPipe | (Delaglio et al., 1995) |  |
| AutoAssign | (Zimmerman et al., 1997, Moseley et al., 2001) | https://montelionelab.chem.rpi.edu/index.htm |
| I-PINE | (Bahrami et al., 2009) |  |
| TALOS+ | (Shen and Bax, 2013) |  |
| *AutoStructure with DP Score (ASDP)* | (Huang et al., 2005b, Huang et al., 2006) | https://montelionelab.chem.rpi.edu/index.htm |
| *Assignment Validation Software (AVS)* | (Moseley et al., 2004) |  |
| *Protein Structure Validation Software Suite (PSVS)* | (Bhattacharya et al., 2007, Moseley et al., 2004) | https://montelionelab.chem.rpi.edu/index.htm |
| MolProbity | (Lovell et al., 2003) |  |
| Procheck | (Laskowski R.A., 1993) |  |
| Prosall | (Sippl, 1993) |  |
| Verify3D | (Lovell et al., 2003) |  |
| PDBStat | (Tejero et al., 2013) |  |
| RPF | (Huang et al., 2005b) | https://montelionelab.chem.rpi.edu/index.htm |
| Matlab |  | Mathworks, https://www.mathworks.com/help/optim/ug/fminunc.html |

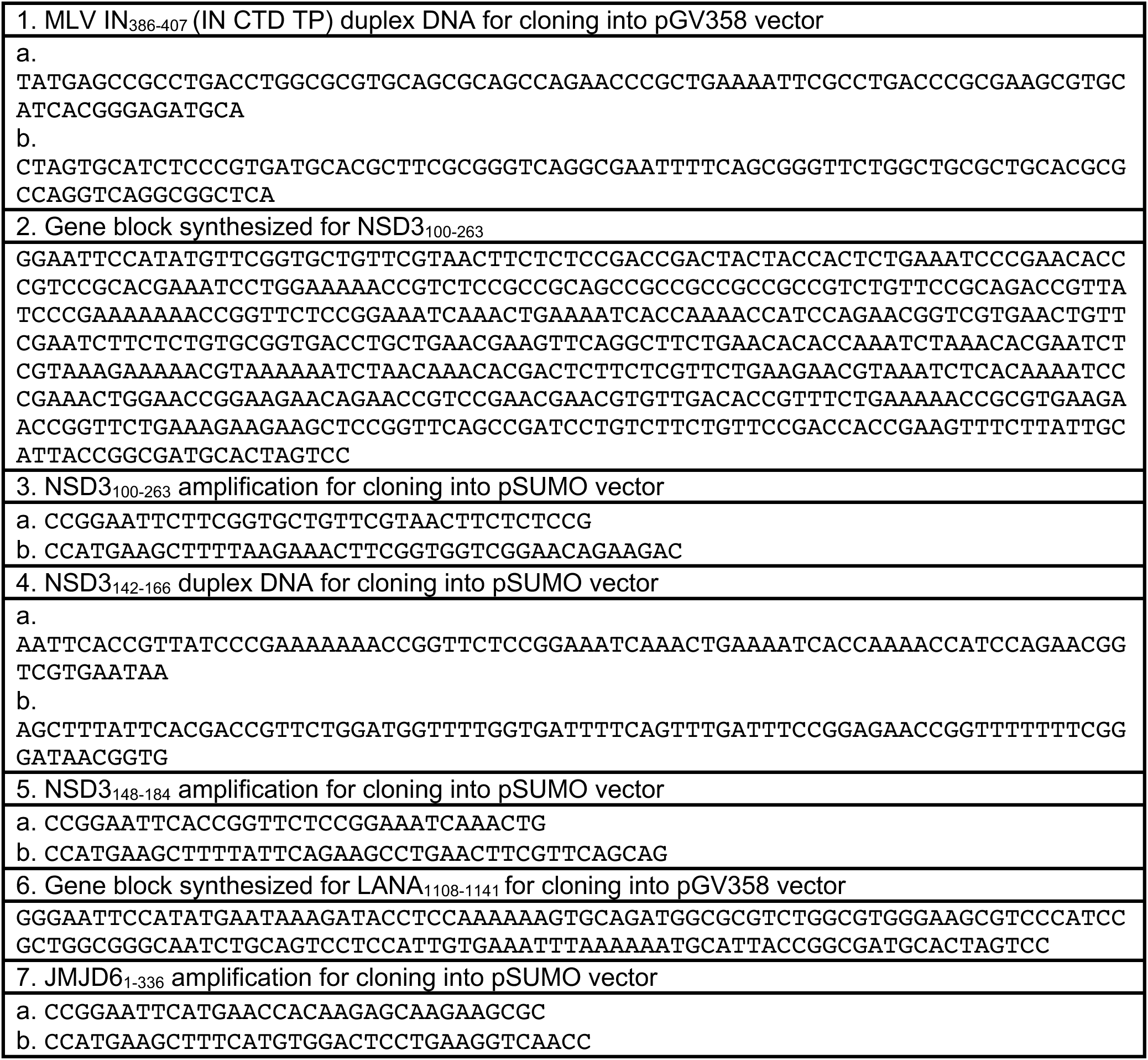
Star Methods Table SM2. List of Oligonucleotides

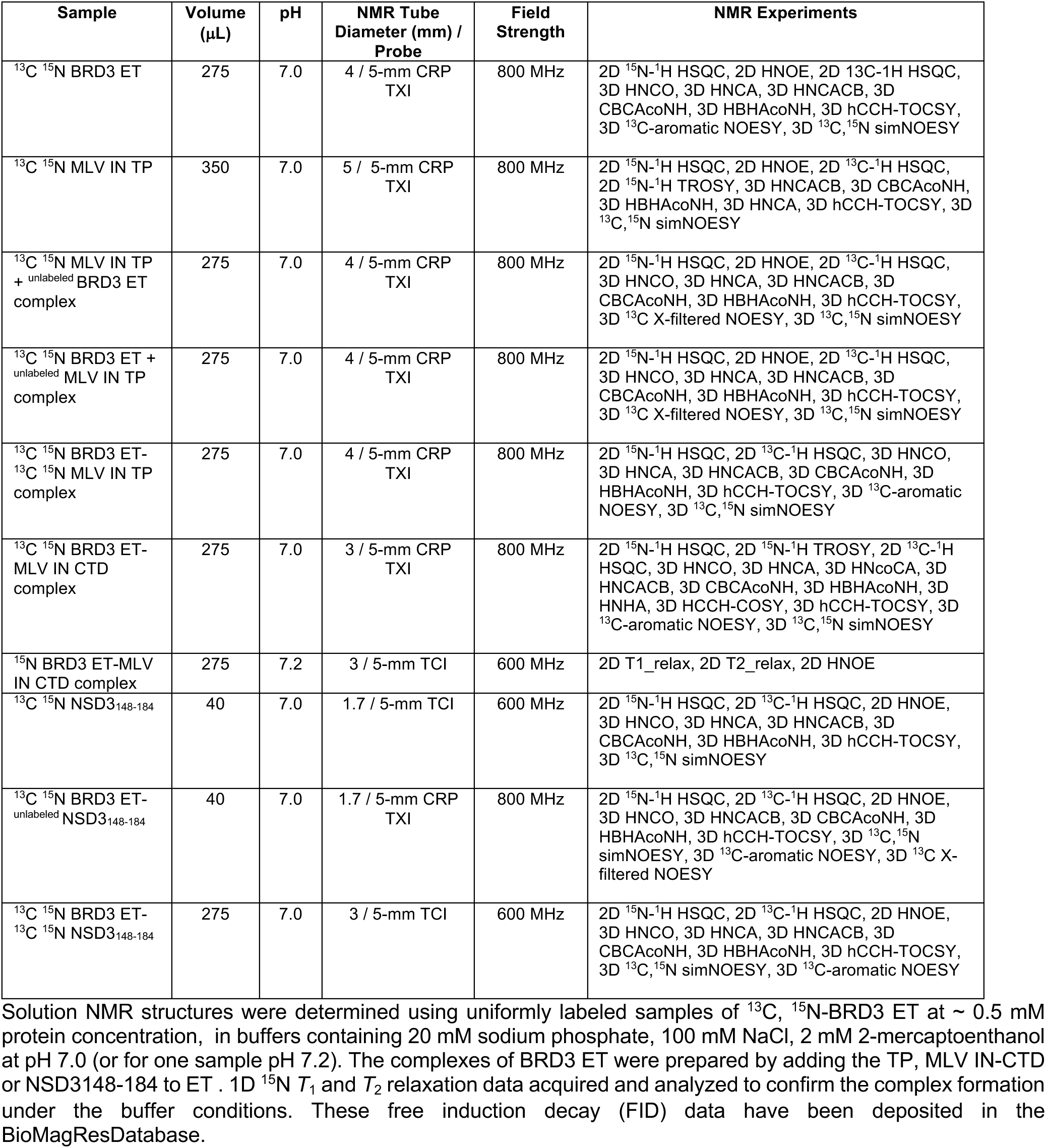
Star Methods Table SM3. NMR samples and data used for resonance assignments, structure determination, and nuclear relaxation measurements.

### Resource Availability

#### Lead Contact

Further information and requests for resources and reagents should be directed to and will be fulfilled by the designated contacts Monica J Roth and Gaetano T. Montelione.

#### Materials Availability

Materials generated in this study are available upon request.

#### Data and Code Availability

BMRB and PDB accession numbers for the structures reported in this paper are: BRD3 ET PDB: 7JMY, BMRB: 30782; BRD 3ET : MLV IN TP PDB: 7JQ8, BMRB: 30786; BRD3 ET : MLV IN CTD PDB: 7JYZ, BMRB: 30791; BRD3 ET : NSD3 PDB: 7JYN, BMRB: 30790.

### Method Details

#### Expression constructs and peptides

Construction of pET15NESG MLV IN CTD and pET15NESG BRD3-ET domain has been described elsewhere (Aiyer et al., 2014). pGV358 (gift from Kushol Gupta, Univ. of Pennsylvania) is a pETDuet based vector that has a C-terminal *Mycobacterium xenopi* GyrA intein (mxe) in-frame followed by the chitin binding domain (CBD) and non-cleavable hexahistidine tag (6xHis). MLV IN CTD TP was generated as follows. Codon optimized complementary oligonucleotides (IDT) encoding for 22 amino acids of the native MLV IN CTD TP sequence and part of the mxe sequence was obtained with overhangs mimicking NdeI and SpeI restriction sites (Star Methods Table SM2). The resulting vector has an open reading frame of M<u>SRLTWRVQRSQNPLKIRLTREA</u>*CITGDAL* (MLV TP is underlined and mxe is italicized), which is in-frame with the rest of the mxe-CBD-6xHis construct. The oligonucleotides were annealed by heating to 94⁰C for 5 mins and slow cooling to 4⁰C for the duration of 1.5 hours. The annealed oligonucleotides were cloned by ligation into pGV358 plasmid digested with NdeI and SpeI. Construction of the CTD of LANA_1108-1141_ into the pGV358 vector was performed using a gene block that was synthesized with NdeI and SpeI sites (Star Methods Table SM2). The corresponding DNA sequences of NSD3_100-263_, NSD3_142-166_, NSD3_148-184_ and JMJD6_1-336_ sequences were encoded in pSUMO fusion vector (Wen et al., 2016) (gift from Volker Vogt, Cornell University). NSD3_100-263_ and NSD3_148-184_ cloning was facilitated by synthesizing a gene block fragment (Star Methods Table SM2) which was amplified through PCR (IDT). JMJD6_1-336_ was amplified using 100 ng of a human cDNA library from 143B cells (Star Methods Table SM2). The ORF of JMJD6_1-336_ contains an EcoRI site, therefore, the cloning was a two-step process involving ligation of an EcoRI-HindIII fragment followed by inserting of an EcoRI-EcoRI fragment. Sanger sequencing was performed to ensure the correct ORF sequences were present in all the plasmids used for expression studies. Unlabeled MLV IN CTD TP, NSD3_148-184_ and LANA_1108-1141_ peptides were procured from Peptide 2.0.

#### Purification of recombinant MLV IN CTD_329-408_ and BRD3 ET_554-640_

Expression and purification of MLV IN CTD and BRD3 ET domain has been described elsewhere (Aiyer et al., 2014). Briefly, induction was carried out in LB media or MJ9 media (Jansson et al., 1996, Aiyer et al., 2014). For isotopically-enriched samples, ^15^NH_4_Cl and ^13^C glucose were the soles sources of nitrogen and carbon, respectively. The bacteria were harvested by centrifugation and resuspended either in lysis buffer [(50 mM Tris-HCl pH 7.5, 500 mM NaCl, 40 mM imidazole, and 1 mM Tris-(2-carboxyethyl)phosphine) or (50 mM sodium phosphate, pH 8.0, 300 mM NaCl, 10 mM imidazole, 10 mM CHAPS, 5 mM 2-mercaptoethanol)], followed by mild sonication. Following Ni-NTA resin purification (Qiagen), fractions eluted in 400 mM imidazole were pooled and concentrated to a volume of less than 250 μl using an Amicon Ultracel-3K centrifugal filter unit (Millipore) as described (Jansson et al., 1996, Aiyer et al., 2014), with the following modifications. The concentrated protein fraction was then injected into an AKTA FPLC and resolved on a Superdex 75 gel filtration column (GE Healthcare) in the buffer for NMR analysis (20 mM sodium phosphate pH 7.0, 100 mM NaCl and 2 mM 2-mercaptoethanol). The eluted fractions were then pooled and concentrated using an Amicon Ultracel-3K centrifugal filter unit (Millipore). These U-^15^N,^13^C-enriched protein samples were stored in this same buffer for NMR analysis.

#### Purification of isotopically-enriched MLV IN CTD TP_386-407_ and NSD3_148-184_

Expression and purification of MLV IN CTD TP_386-407_-mxe-CBD-6xHis fusion protein construct from the pGV358 backbone was adapted from previous studies (Aiyer et al., 2015, Aiyer et al., 2014, Schneider et al., 2012), resulting in the IN CTP TP fused to the self-cleaving intein-Chitin binding domain (CBD). Induction for protein expression was carried out in MJ9 media (Jansson et al., 1996), in which ^15^NH_4_Cl and uniformly ^13^C-enriched glucose are the sole sources of nitrogen and carbon, respectively. The bacteria were harvested by centrifugation and resuspended either in lysis buffer [50 mM Tris-HCl pH 7.5, 500 mM NaCl, 40 mM imidazole, and 1 mM Tris-(2-carboxyethyl)phosphine] or (50 mM sodium phosphate, pH 8.0, 300 mM NaCl, 10 mM imidazole, 10 mM CHAPS, 5 mM 2-mercaptoethanol)], followed by mild sonication. The IN TP-CBD fusion protein was purified first using NiNTA beads (Qiagen) and then bound to chitin beads (New England Biolabs) as per the respective manufacturer’s recommendations. Isotopically enriched peptide was released from the fusion protein bound to chitin agarose beads (NEB) by incubating for 48 hours at room temperature with 20 mM sodium phosphate pH 8.0, 0.2 M NaCl, 0.1 mM EDTA, 75 mM dithiothreitol (DTT) (Fluka) to initiate self-cleaving reaction under reducing conditions resulting in on-column cleavage. Unbound cleaved peptide was then separated from uncleaved fusion protein and mxe-CBD-6xHis by passing the suspension through buffer equilibrated (without DTT) fresh chitin resin (NEB) settled in a gravimetric econopack column (BioRad). Purified isotopically enriched peptide was concentrated and buffer exchanged to 20 mM sodium phosphate pH 7.0, 100 mM NaCl and 2 mM DTT using 3 kDa MWCO Ultragel Amicon filters (Millipore) or 2 kDa MWVO Hydrosart filters (Vivaspin). Homogeneity (> 95%) was validated by SDS-PAGE, and isotope-enrichment by MALDI-TOF mass spectrometry (Supplementary Figure S2).

Isotope-enriched NSD3_148-184_ was produced as a SUMO-fusion construct in *E. coli* expression hosts and cleaved with SUMO protease. Bacteria were harvested by centrifugation and resuspended in lysis buffer [50 mM Tris-HCl pH 7.5, 500 mM NaCl, 40 mM imidazole, and 1 mM Tris-(2-carboxyethyl)phosphine], followed by mild sonication. After the removal of cell debris by centrifugation at 27,000 × g for 30 min at 4 °C, the supernatant was applied to a 5 ml HisTrap^TM^ affinity column (GE Healthcare). The column was washed with the lysis buffer and the protein was eluted using a buffer containing 50 mM Tris-HCl pH 7.5, 500 mM NaCl, 300 mM imidazole, and 1 mM Tris-(2 carboxyethyl) phosphine. Subsequently, the eluted protein sample, containing the SUMO fusion of NSD3_148-184_, was further purified by gel filtration (HiLoad 16/60 Superdex 75; GE Healthcare) chromatography in a cleavage buffer (50 mM Tris-HCl pH 8.0, 100 mM NaCl, and 10 mM DTT). The cleavage of the fusion protein was carried out by adding an aliquot (1:50–100 mass ratio) of yeast SUMO protease Ulp1 containing an N-terminal 6xHis tag (Ii et al., 2007), and the sample was then incubated at 25 °C for 18-24 hours. The degree of the cleavage was monitored by SDS-PAGE. To remove the uncleaved fusion protein, SUMO, and SUMO protease, the mixture was applied to a 5 ml HisTrap ^TM^ column. The flow through and wash fractions, which contain the peptide NSD3_148-184_ were verified by SDS-PAGE, and pooled. The purified NSD3_148-184_ was > 95% pure as determined by SDS-PAGE (Supplementary Figure S2).

#### NMR Assignments and Structure Determination

^13^C,^15^N-enriched samples of BRD3 ET and its complexes with MLV IN TP, MLV IN CTD and NSD3_148-184_ at pH 7.0 were used for structure determination. These samples are summarized in Star Methods Table SM3. Samples were prepared in 3-, 4- or 5-mm Shigemi NMR tubs, or in 1.7-mm micro NMR tubes. Conventional 3D triple resonance (Baran et al., 2004, Huang et al., 2005a) and 3D NOESY NMR experiments were executed using standard Bruker NMR pulse sequences optimized for each protein sample at 25 °C on Bruker Avance III 600 MHz and Avance III 800 MHz NMR spectrometer systems. Complex formation was confirmed using 1D ^15^N T_1_ and T_2_ relaxation experiments to estimate rotational correlation times τ_c_, and effective molecular weights were estimated by comparison with τ_c_ measurements made on collection of reference proteins, as described elsewhere (Rossi et al., 2010). 1D ^15^N T_1_ and T_2_ relaxation data was processed using Bruker Topspin 2.3 software and analyzed using the “t1guide” module in Topspin to obtain ^15^N T_1_ and T_2_ values. All NMR data was processed using *NMRPipe* (Delaglio et al., 1995) and visualized using *NMRFaM SPARKY* (Lee et al., 2015).

Initial backbone ^1^H, ^13^C, and ^15^N resonance assignments were determined automatically using both the I-Pine server (Bahrami et al., 2009, Lee et al., 2019) and *AutoAssign* (Zimmerman et al., 1997, Moseley et al., 2001) software, and then refined by interactive manual analysis with the program *Sparky* (Lee et al., 2015). Side chain resonance assignments were completed manually using *Sparky*. The NMR experiments used to determine these resonance assignments for each free ET and each complex are summarized in Star Methods Table SM3. NMR resonance assignment validation was done using the program Assignment Validation Software (AVS) (Moseley et al., 2004).

NOESY data collection included 3D N-edited-NOESY-HSQC, 3D C-edited NOESY-HSQC, and 3D sim N,C-edited-HSQC (Cavanagh et al., 2006), and well as 3D ^13^C X-filtered NOESY experiments (Stuart et al., 1999) for detecting intermolecular NOE interactions between labeled and unlabeled fragments. NOESY mixing times were 100 to 120 ms. Peak intensities from a 3D NOESY spectra, together with broad dihedral angle constraints determined from backbone chemical shift data using TALOS+ (Shen and Bax, 2013) for ordered residues with TALOS+ confidence scores of 10, were used as input for structure calculations. NOESY peak assignments were made automatically using the *CANDID* module of *Cyana* (Guntert et al., 1997, Herrmann et al., 2002) together with the program *ASDP* (Huang et al., 2005b, Huang et al., 2006). The resulting structures were then refined in explicit water using CNS (Brunger et al., 1998).

Structure quality analyses were performed using the *Protein Structure Validation Suite* (PSVS) (Bhattacharya et al., 2007, Moseley et al., 2004), which includes the *MolProbity* (Lovell et al., 2003), *Procheck* (Laskowski R.A., 1993), *Verify3D* (Lovell et al., 2003) and *ProsaII* (Sippl, 1993), *RPF* (Huang et al., 2005b) and *PDBStat* (Tejero et al., 2013) software packages. The final ensemble of 20 models (excluding the C-terminal His_6_ tag), NMR resonance assignments, and extensive raw NMR fid data and spectral peak lists were deposited into the Protein Data Bank and BioMagRes Database.

#### ^15^N Nuclear Relaxation Studies of IN CTD : ET Complex

^15^N nuclear relaxation experiments were performed on ^15^N-enriched sample of the IN CTD:ET complex in 20 mM sodium phosphate, 100 mM NaCl and 2 mM 2-mercaptoethanol, at pH 7.2. A series of 2D NMR spectra were acquired as described elsewhere (Feng et al., 1998). Data collection for T_1_ used decays delays of 20, 50, 100, 300, 500, 700, 900, 1200 and 2000 ms, and T*_2_* measurements were made with decay delays of 17, 34, 51, 68, 85, 96, 119, 170 and 220 ms. ^1^H-^15^N steady-state NOE values were measured with two different data sets, one collected with no initial proton saturation and a second with initial proton saturation of 3 s. These 2D NMR spectra were processed and phased identically using *NMRPIPE*, and peak intensities obtained using SPARKY software. Steady heteronuclear ^15^N-^1^H NOE values were obtained from the ratios of cross peak intensities in the saturated spectrum to those in the unsaturated spectrum (I_sat_ / I_eq_).

#### Error Estimates in Nuclear Relaxation Measurements

Curve fits to exponential decay curves for estimating transverse (T_1_) and longitudinal (T_2_) ^15^N relaxation rates were made using a global optimization function. Errors were estimated by Gaussian probability distribution modeling, using the standard deviation of the measured intensity uncertainties (± 5%) about each intensity in the data.

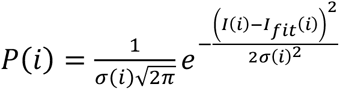

The sum over P(i) from all time series data points i in the time series is maximized to find the global best fit *I_fit_*(*i*) to all the data through the measured intensities *I*(*i*). This analysis accounts for the experimental measurement uncertainty, defined by σ(i), of ± 5%. This approach results in the best fit of data to the single-exponential decay model, i.e. *I_fit_*(*i*) = *a* * *exp*(−*x*/*b*) + *c*, given the uncertainty in the intensity values I(i) of σ(i), and maximizes the exponential fit, in particular the decay constants T_1_ and T_2_, within the one sigma bounds of the measurement uncertainties. The root mean square error of this fit was used to find the largest and smallest decay constants about this most likely fit, giving the fit uncertainties from the estimate of the possible measurement errors. The correlation time τ_c_ for each amide N-H bond vector, and its uncertainty, were then determined by finding an exact solution to the ratio of the contributing spectral densities which define T_1_/T_2_ (Carper and Keller, 1997). These T_1_, T_2_, and τ_c_ calculations and their fitted uncertainties used the minimization algorithm fminunc in *Matlab* (Mathworks). Heteronuclear NOE (HNOE) values (I_sat_/I_eq_) and their uncertainties, σ_HNOE_, were determined as described elsewhere (Feng et al., 1998), assuming a standard deviation (σ_i_) of the measured resonance intensities I(i) of ± 5%.

#### Chemical shift perturbations measurements

The threshold for defining a significant chemical shift perturbation (CSP) was determined from the following iterative analysis, as described elsewhere (Ma et al., 2016). The standard deviation (σ) of the shift changes was first calculated. To prevent biasing the distribution by including the small number of residues with very large shift changes, any residues for which the shift change is greater than 3σ were excluded. The next standard deviation was then recalculated excluding residues with shift changes more than 3σ. Iteration of these calculations was performed until no further residues are excluded, resulting in deviation for unperturbed sites of 3σ = 0.02 ppm. Accordingly, we conservatively choose 0.05 ppm (50 ppb) as the threshold to minimize any false positives.

---

**Supplementary Table S1.**
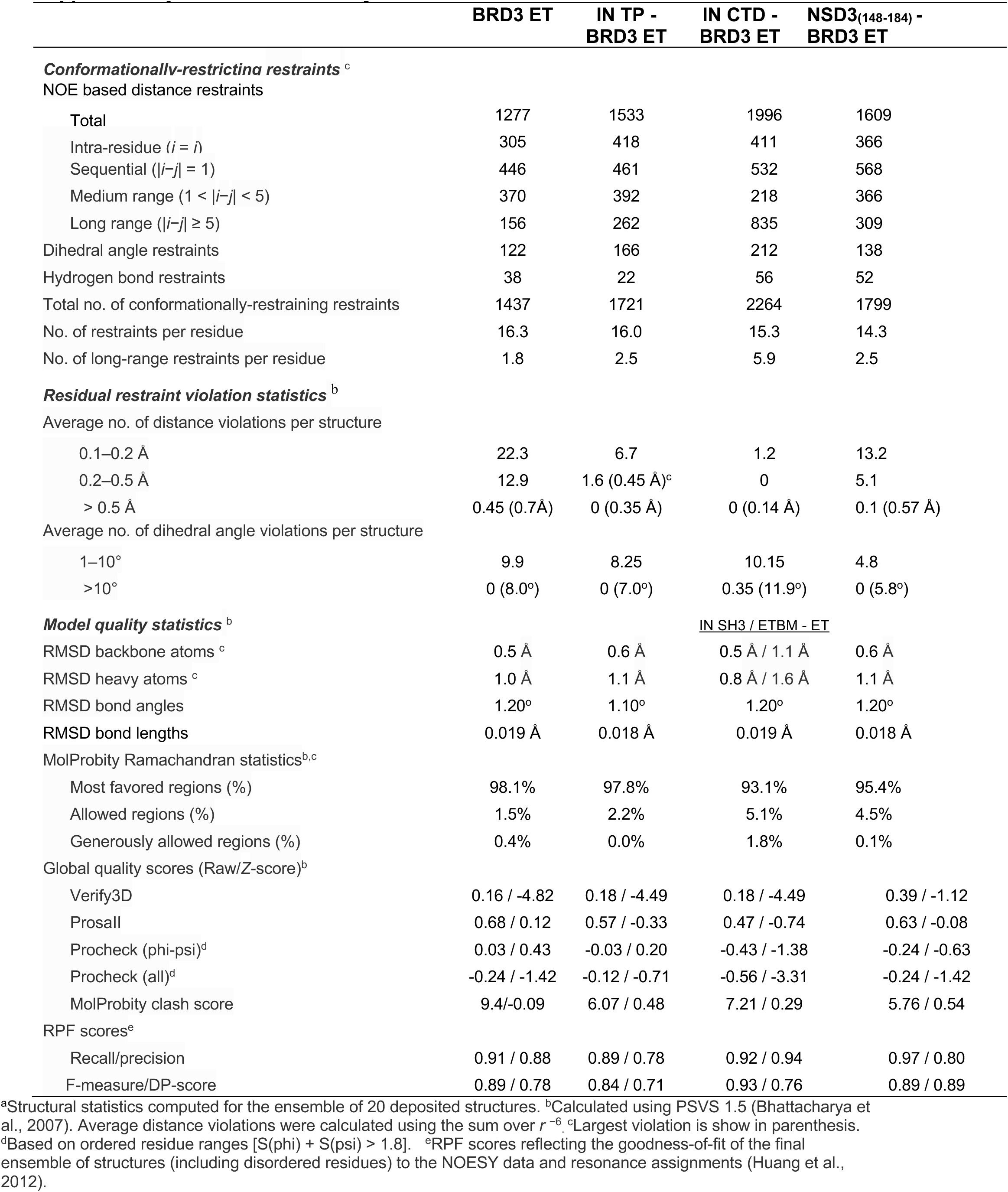
Summary of structural statistics^a^

|  | BRD3 ET | IN TP -<br>BRD3 ET | IN CTD -<br>BRD3 ET | NSD3<sub>(148-184)</sub> -<br>BRD3 ET |
| --- | --- | --- | --- | --- |
| **Conformationally-restricting restraints<sup>c</sup>** |  |  |  |  |
| NOE based distance restraints |  |  |  |  |
| Total | 1277 | 1533 | 1996 | 1609 |
| Intra-residue ($i = j$) | 305 | 418 | 411 | 366 |
| Sequential ($\vert i-j\vert = 1$) | 446 | 461 | 532 | 568 |
| Medium range ($1 < \vert i-j\vert < 5$) | 370 | 392 | 218 | 366 |
| Long range ($\vert i-j\vert \geq 5$) | 156 | 262 | 835 | 309 |
| Dihedral angle restraints | 122 | 166 | 212 | 138 |
| Hydrogen bond restraints | 38 | 22 | 56 | 52 |
| Total no. of conformationally-restraining restraints | 1437 | 1721 | 2264 | 1799 |
| No. of restraints per residue | 16.3 | 16.0 | 15.3 | 14.3 |
| No. of long-range restraints per residue | 1.8 | 2.5 | 5.9 | 2.5 |
| **Residual restraint violation statistics<sup>b</sup>** |  |  |  |  |
| Average no. of distance violations per structure |  |  |  |  |
| 0.1–0.2 Å | 22.3 | 6.7 | 1.2 | 13.2 |
| 0.2–0.5 Å | 12.9 | 1.6 (0.45 Å)<sup>c</sup> | 0 | 5.1 |
| > 0.5 Å | 0.45 (0.7 Å) | 0 (0.35 Å) | 0 (0.14 Å) | 0.1 (0.57 Å) |
| Average no. of dihedral angle violations per structure |  |  |  |  |
| 1–10° | 9.9 | 8.25 | 10.15 | 4.8 |
| >10° | 0 (8.0°) | 0 (7.0°) | 0.35 (11.9°) | 0 (5.8°) |
| **Model quality statistics<sup>b</sup>** |  |  | <u>IN SH3 / ETBM - ET</u> |  |
| RMSD backbone atoms<sup>c</sup> | 0.5 Å | 0.6 Å | 0.5 Å / 1.1 Å | 0.6 Å |
| RMSD heavy atoms<sup>c</sup> | 1.0 Å | 1.1 Å | 0.8 Å / 1.6 Å | 1.1 Å |
| RMSD bond angles | 1.20° | 1.10° | 1.20° | 1.20° |
| RMSD bond lengths | 0.019 Å | 0.018 Å | 0.019 Å | 0.018 Å |
| MolProbity Ramachandran statistics<sup>b,c</sup> |  |  |  |  |
| Most favored regions (%) | 98.1% | 97.8% | 93.1% | 95.4% |
| Allowed regions (%) | 1.5% | 2.2% | 5.1% | 4.5% |
| Generously allowed regions (%) | 0.4% | 0.0% | 1.8% | 0.1% |
| Global quality scores (Raw/Z-score)<sup>b</sup> |  |  |  |  |
| Verify3D | 0.16 / -4.82 | 0.18 / -4.49 | 0.18 / -4.49 | 0.39 / -1.12 |
| ProsaII | 0.68 / 0.12 | 0.57 / -0.33 | 0.47 / -0.74 | 0.63 / -0.08 |
| Procheck (phi-psi)<sup>d</sup> | 0.03 / 0.43 | -0.03 / 0.20 | -0.43 / -1.38 | -0.24 / -0.63 |
| Procheck (all)<sup>d</sup> | -0.24 / -1.42 | -0.12 / -0.71 | -0.56 / -3.31 | -0.24 / -1.42 |
| MolProbity clash score | 9.4/-0.09 | 6.07 / 0.48 | 7.21 / 0.29 | 5.76 / 0.54 |
| RPF scores<sup>e</sup> |  |  |  |  |
| Recall/precision | 0.91 / 0.88 | 0.89 / 0.78 | 0.92 / 0.94 | 0.97 / 0.80 |
| F-measure/DP-score | 0.89 / 0.78 | 0.84 / 0.71 | 0.93 / 0.76 | 0.89 / 0.89 |

<sup>a</sup>Structural statistics computed for the ensemble of 20 deposited structures. <sup>b</sup>Calculated using PSVS 1.5 (Bhattacharya et al., 2007). Average distance violations were calculated using the sum over $r^{-6}$ . <sup>c</sup>Largest violation is shown in parenthesis.
<sup>d</sup>Based on ordered residue ranges [S(phi) + S(psi) > 1.8]. <sup>e</sup>RPF scores reflecting the goodness-of-fit of the final ensemble of structures (including disordered residues) to the NOESY data and resonance assignments (Huang et al., 2012).

**Supplementary Fig. S1.**
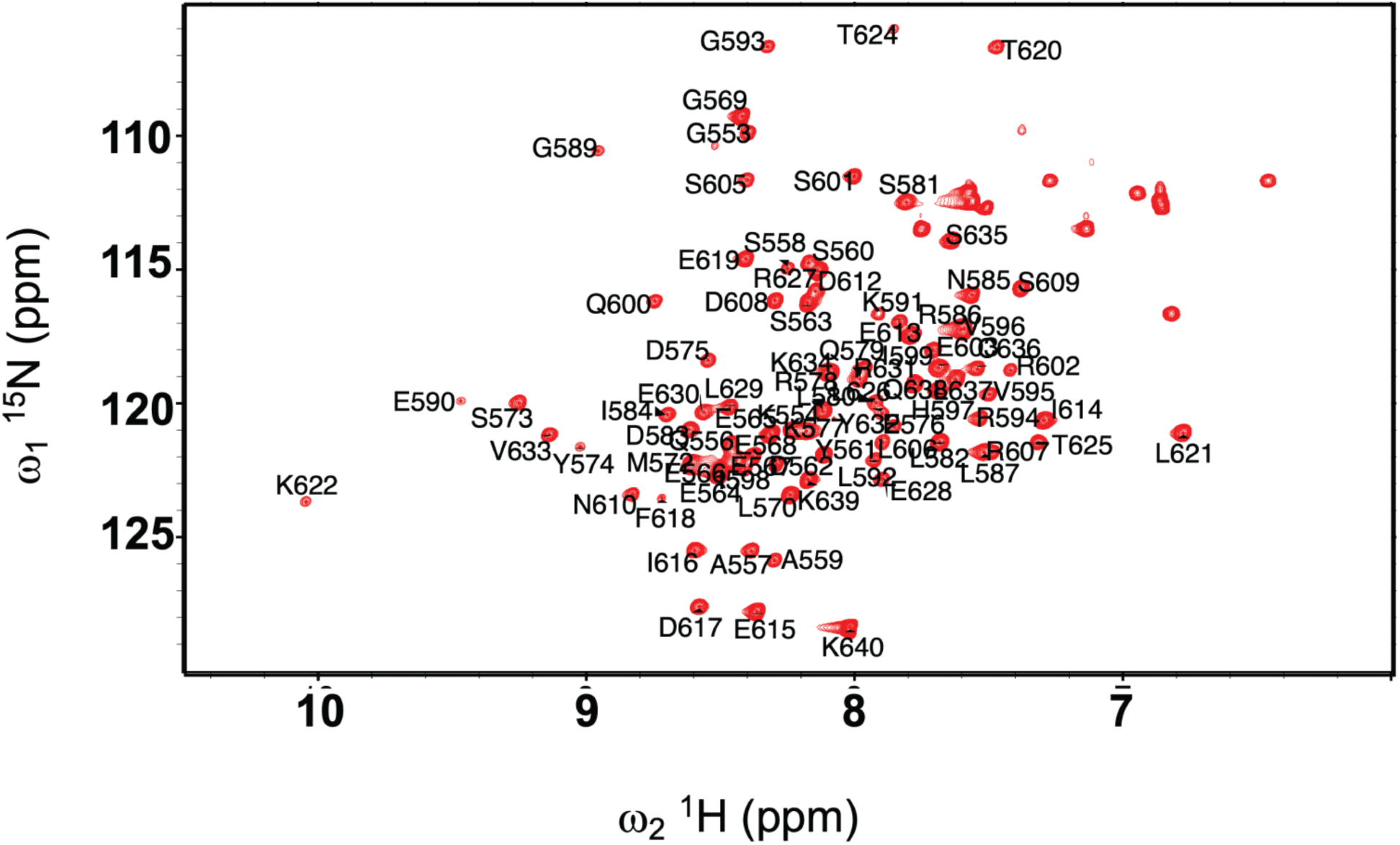
[^15^N-^1^H]-HSQC of BRD3 ET domain. 2D [^15^N,^1^H]-HSQC spectrum of ^13^C,^15^N BRD3 ET acquired at 25 °C using 45 μL of 0.5 mM sample at pH 7.0 and 600 Hz Bruker Avance NMR spectrometer equipped with a 1.7-mm micro cryoprobe. Backbone amide ^15^N, ^1^H^N^ resonance assignments are labeled in black.

**Supplementary Fig. S2.**
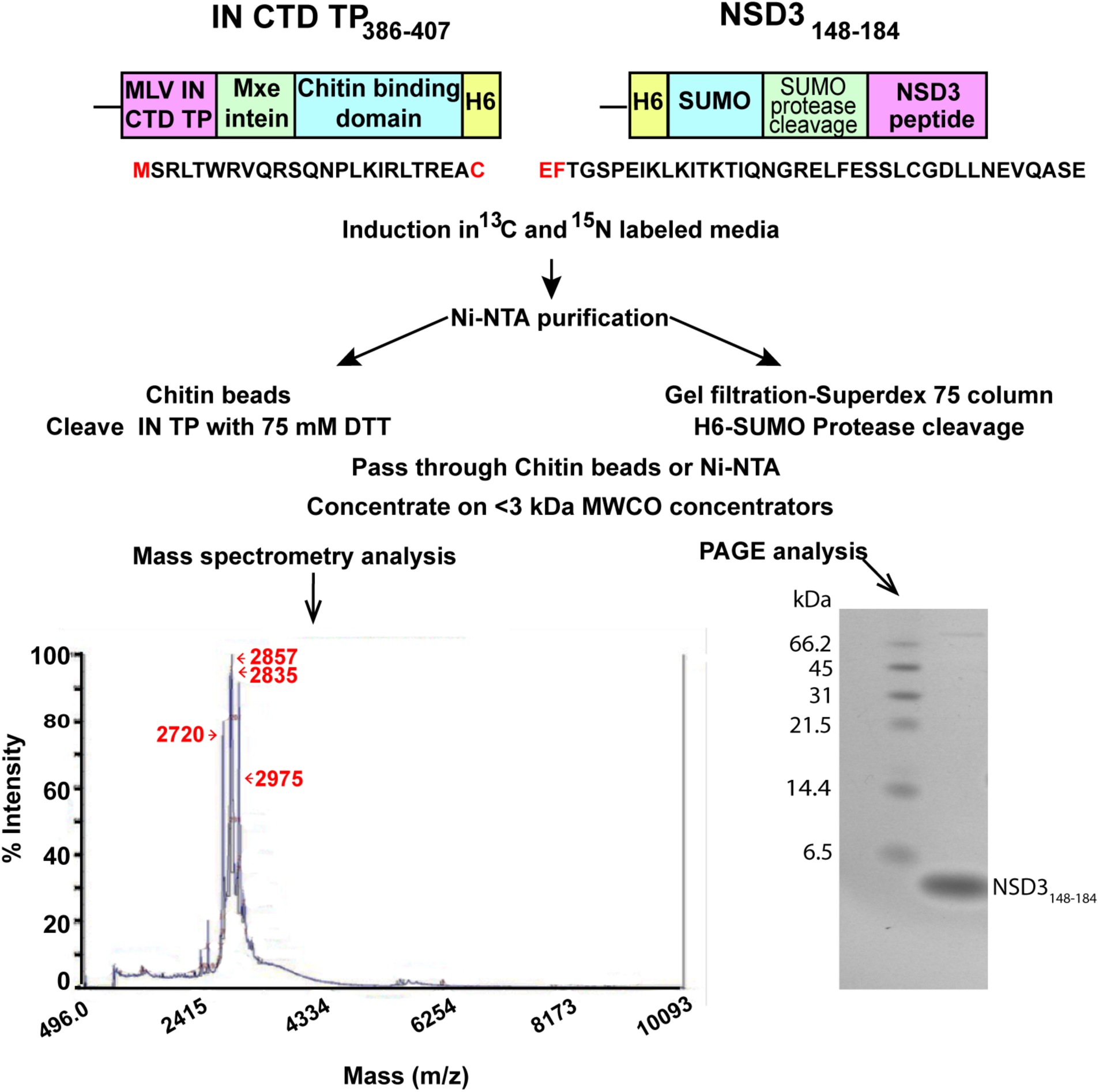
Peptide purification schematic. The schematic of the IN CTD TP_386-407_ chitin binding domain (CBD) fusion protein (left) and the NSD3_148-184_-SUMO fusion protein (right) are illustrated at the top. Position of the His_6_ tag (H6) is indicated. The sequences of the peptides are indicated below each construct; amino-acid residues indicated in red are not encoded by the peptides of interest. Purification of the peptides follow parallel protocols, involving column chromatography and either chemical or enzymatic cleavage of the tag away from the peptides of interest. Exemplary mass spectrometry of the TP_386-407_ is shown on the left. The predicted molecular mass of the labeled peptide containing the N-terminal Met is 2830. Species smaller migrating at 2720 could result from loss of the terminal Met. Products larger at 2975 Da could result from incomplete removal of the DTT adduct after cleavage of the CBD. Exemplary Coomassie staining of the purified NSD3 peptide is shown on the bottom right. Molecular weight standards are as indicated.

**Supplementary Fig. S3.**
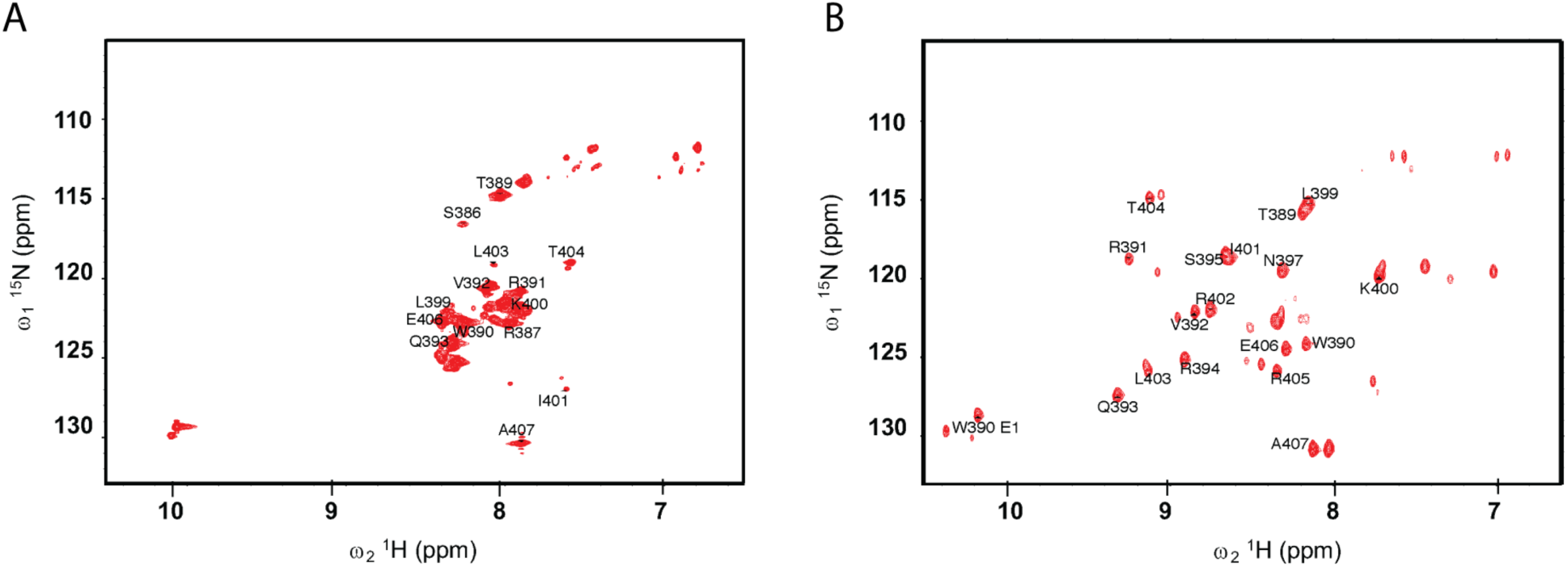
Perturbations for the ^15^N,^13^C-enriched TP without and with ET domain. (A) [^15^N-^1^H]-HSQC spectrum of unbound ^15^N labeled MLV IN CTD TP. (B) [^15^N-^1^H]-HSQC spectrum of ^15^N labeled MLV IN CTD TP bound to unlabeled BRD3 ET domain. Assignments of some resonances of TP are indicated.

**Supplementary Fig. S4.**
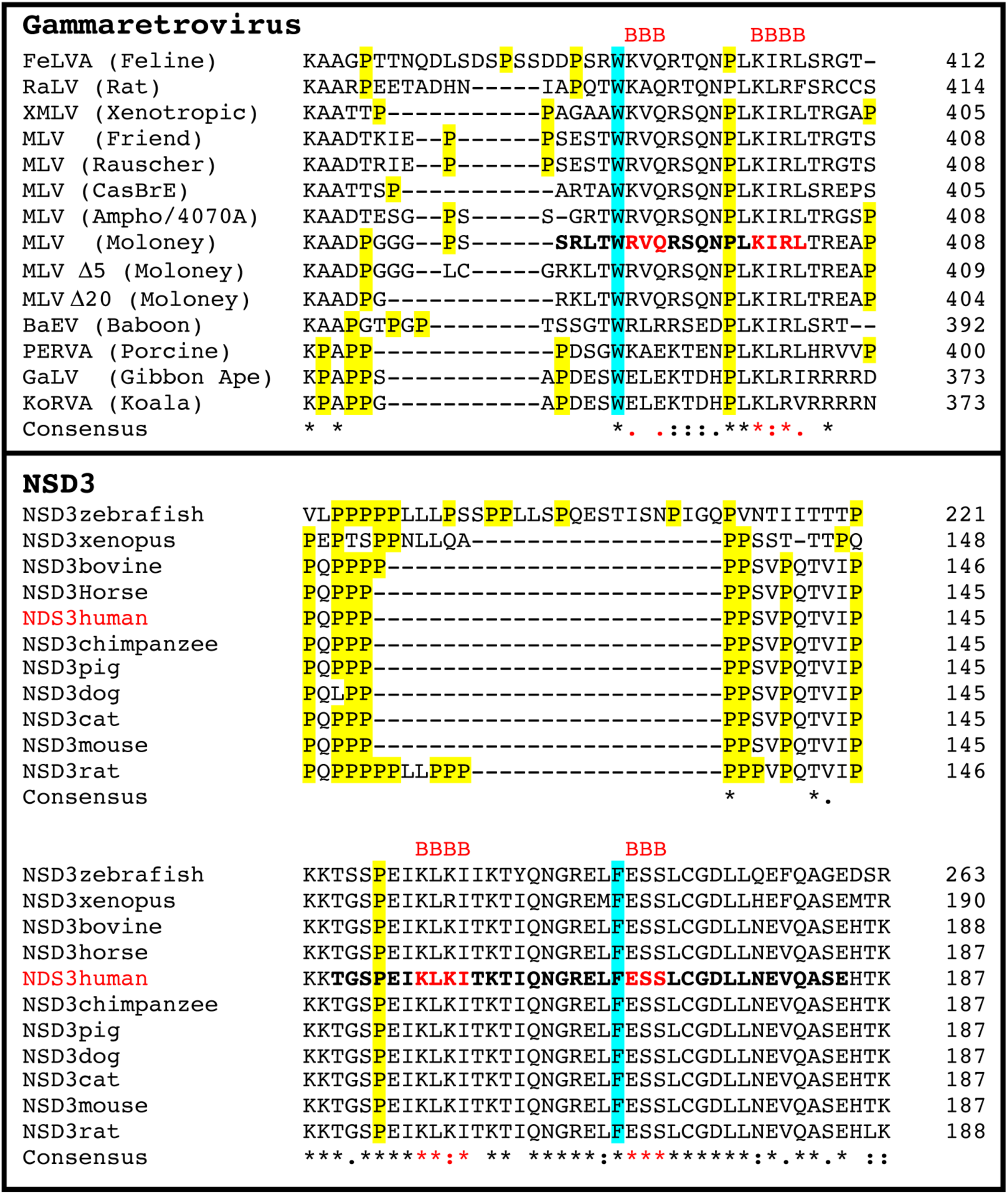
Alignment of gammaretroviral IN linker regions (top) and NSD3 (bottom) proteins. Alignments were performed using Clustal Omega (1.2.4) (Madeira et al., 2019, Sievers et al., 2011, Sievers and Higgins, 2018). Gammaretroviruses analyzed include: P10273- Feline leukemia virus, Q83379- Rat Leukemia virus, MH450110.1- Xenotropic murine leukemia virus (MLV) isolate AKR6, O39735- Friend murine leukemia virus, O41250- Rauscher murine leukemia virus, P08361- Cas-Br-E MLV, ABU54793.1- Amphotropic MLV 4070A, P03355- Moloney murine leukemia virus, P10272- Baboon endogenous virus, Q8J4V8- PERV-A, P21414- GaLV, and Q9TTC1- KoRV. M-MLV Δ5 and Δ 20 were isolated as described (Loyola et al., 2019). NSD3 protein sequences analyzed include: F1QV68- NSD3 (zebrafish), A0A1L8H2H2- NSD3 (xenopus), E1BNH7- NSD3 (bovine), F6WYL3- NSD3 (horse), Q9BZ95-NSD3 (human), H2R3Q8- NSD3 (chimpanzee), A0A286ZK72-NSD3 (pig), E2QUJ0-NSD3 (dog), M3WJ59-NSD3 (cat), Q6P2L6- NSD3 (mouse), and D3ZK47- NSD3 (rat). Consensus sequences notations are: (*) fully conserved residue, (:) conservation between groups of strongly similar properties, and (.) conservation between groups of weakly similar properties (Gonnet et al., 1992). IN-TP and NSD3 peptide used in these studies are indicated in bold. Pro are highlighted in yellow. Conserved Trp for MLV-IN and Phe for NSD3 that form the cap to the hydrophobic network are highlighted in cyan. Position and conservation of the two beta-strands (B) of MLV IN and NSD3 interacting with the ET domain are indicated in red.

**Supplementary Fig. S5.**
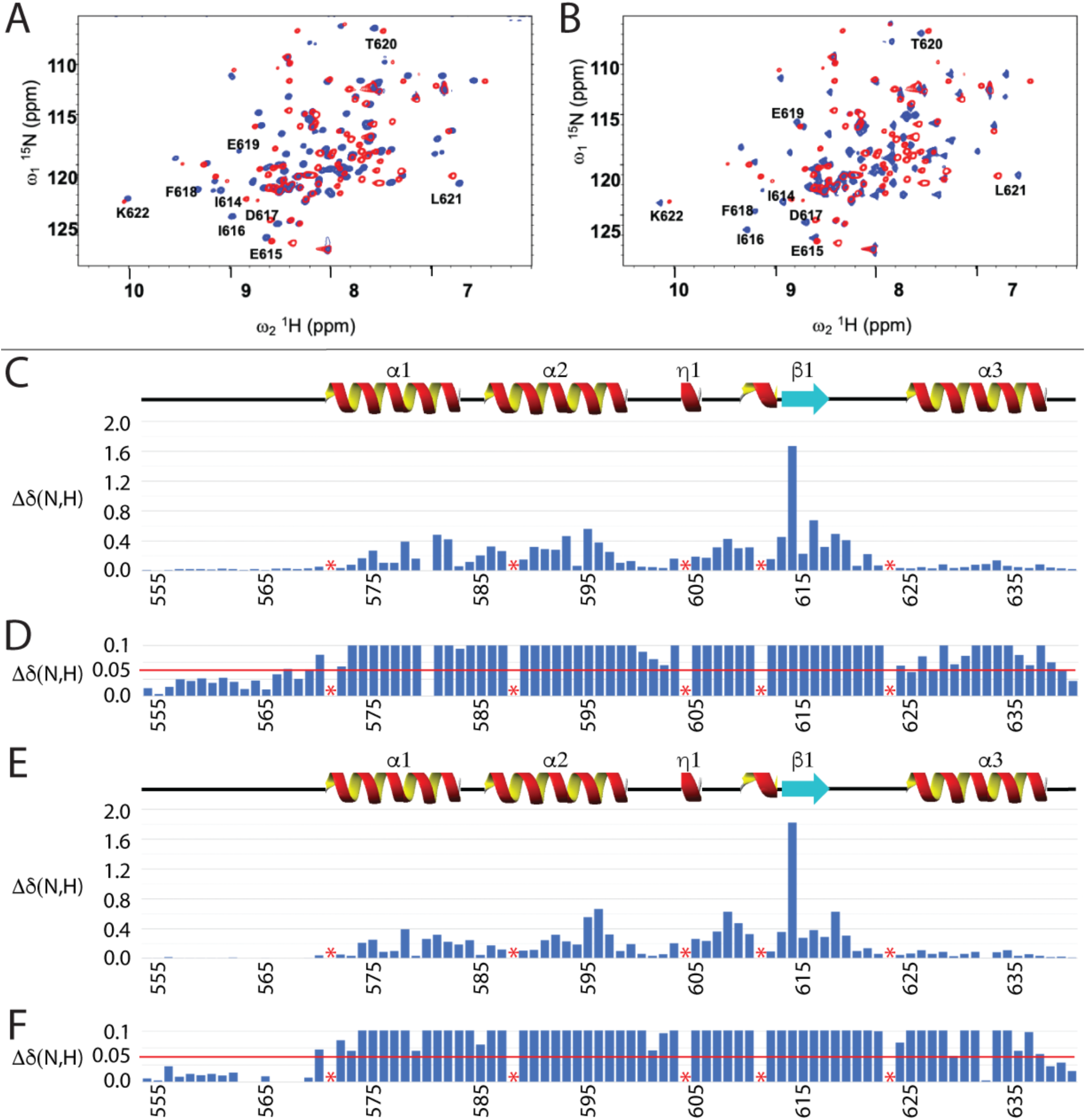
Comparison of chemical shift perturbations of BRD3 ET induced by IN TP and NSD3. Panels A and B: HSQC of the BRD3 ET (blue) in the presence of unlabeled IN TP (panel A, red) and NSD3 (panel B, red). Chemical shift perturbations of BRD3 ET domain relative to the bound (C, D) TP and (E, F) NSD3_148-184_calculated using Δδ(N,H)= ((δN*6)^2^+(δH)^2^)^0.5^ and plotted as a function of BRD3 ET sequence. Panels D and F show an expansion of the data in Panels C and E, demonstrating that most residues have Δδ(N,H) greater than 0.05 ppm (horizontal red lines), a typical cutoff value for unperturbed chemical shift changes on complex formation (Ma et al., 2016). Amino-acid residue positions marked with a red * correspond to Pro. Assignments of some resonances of ET are indicated. Secondary structure of BRD3 ET is shown schematically in panels C and E.

**Supplementary Fig. S6.**
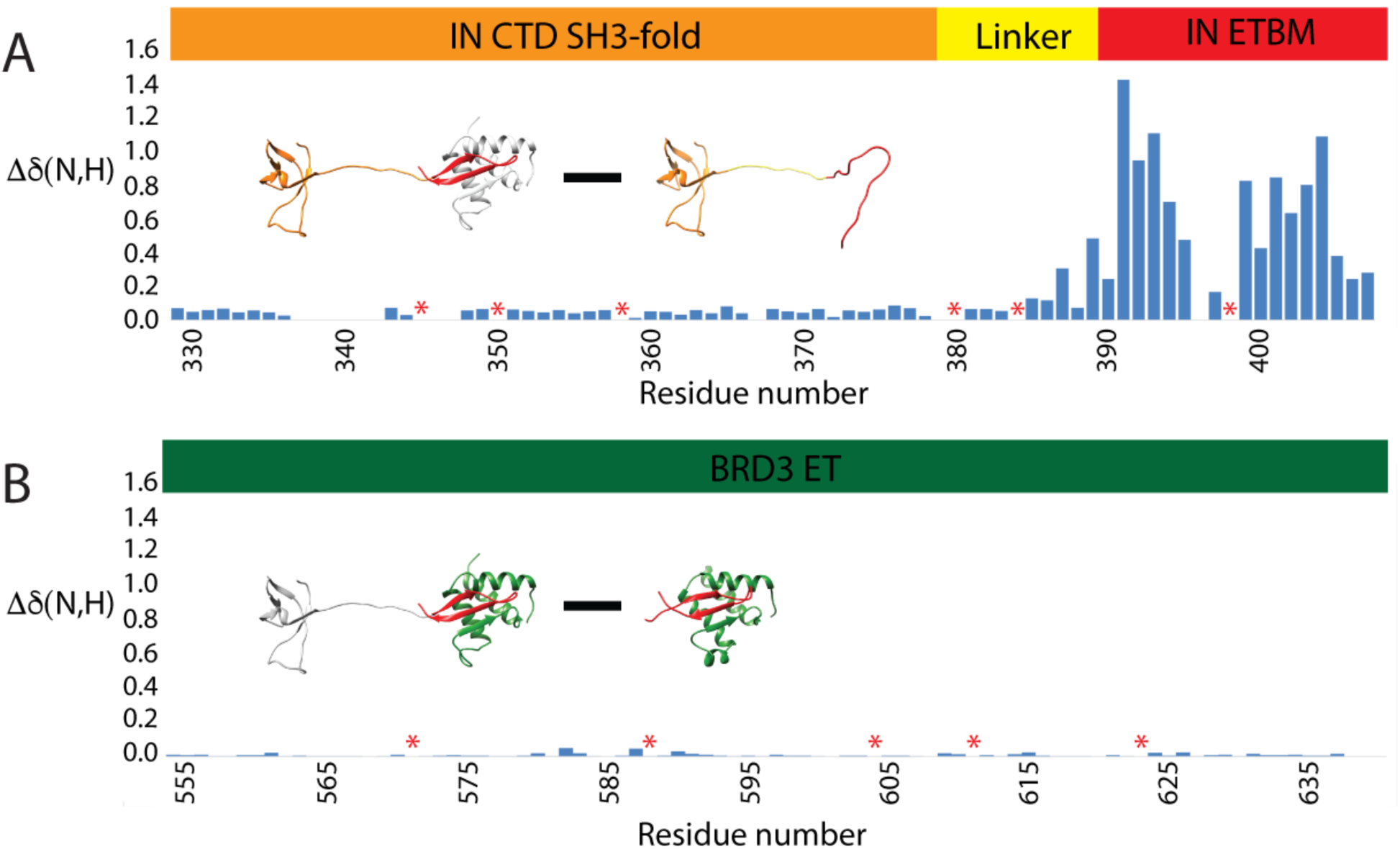
Comparison of chemical shift perturbations of backbone amides. (A) Chemical shift perturbations of MLV IN CTD, in complex with BRD3 ET minus chemical shifts of apo MLV IN CTD. (B) Chemical shift perturbations of BRD3 ET, in complex with MLV IN CTD : BRD3 ET minus chemical shifts of MLV IN TP : BRD3 ET. Inset of the ribbon diagrams in color indicate the amide ^15^N,^1^H resonances being compared while the rest is shown in gray. Residue numbers of the IN CTD and BRD3 ET domain are indicated in each panel (bottom) along with the schematic of the domains (top). Residues indicated by a red * correspond to Pro.

**Supplementary Fig. S7.**
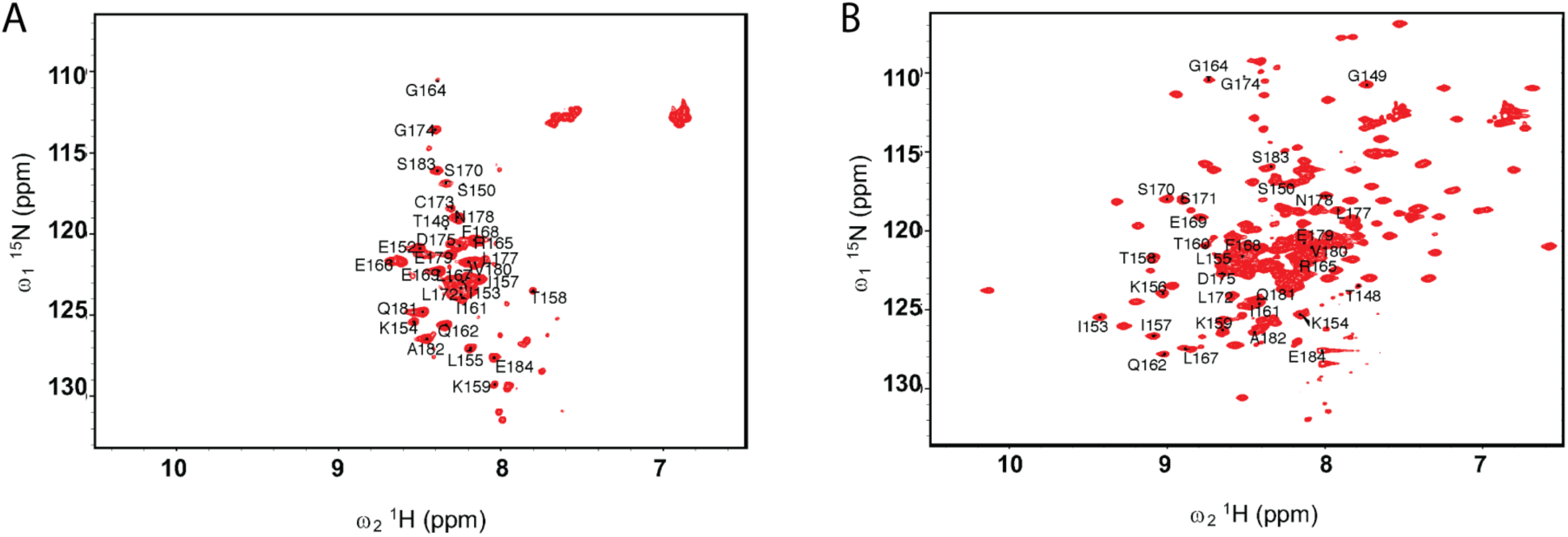
Perturbations for the ^15^N,^13^C-enriched NSD3_148-184_ without and with ET domain. (A) 2D ^15^N-^1^H HSQC spectrum of ^13^C, ^15^N-NSD3_148-184_ and (B) ^13^C, ^15^N-NSD3_148-184_-^13^C, ^15^N-BRD3 ET complex. Assignments of some resonances of NSD3_148-184_ are indicated.

**Supplementary Fig. S8.**
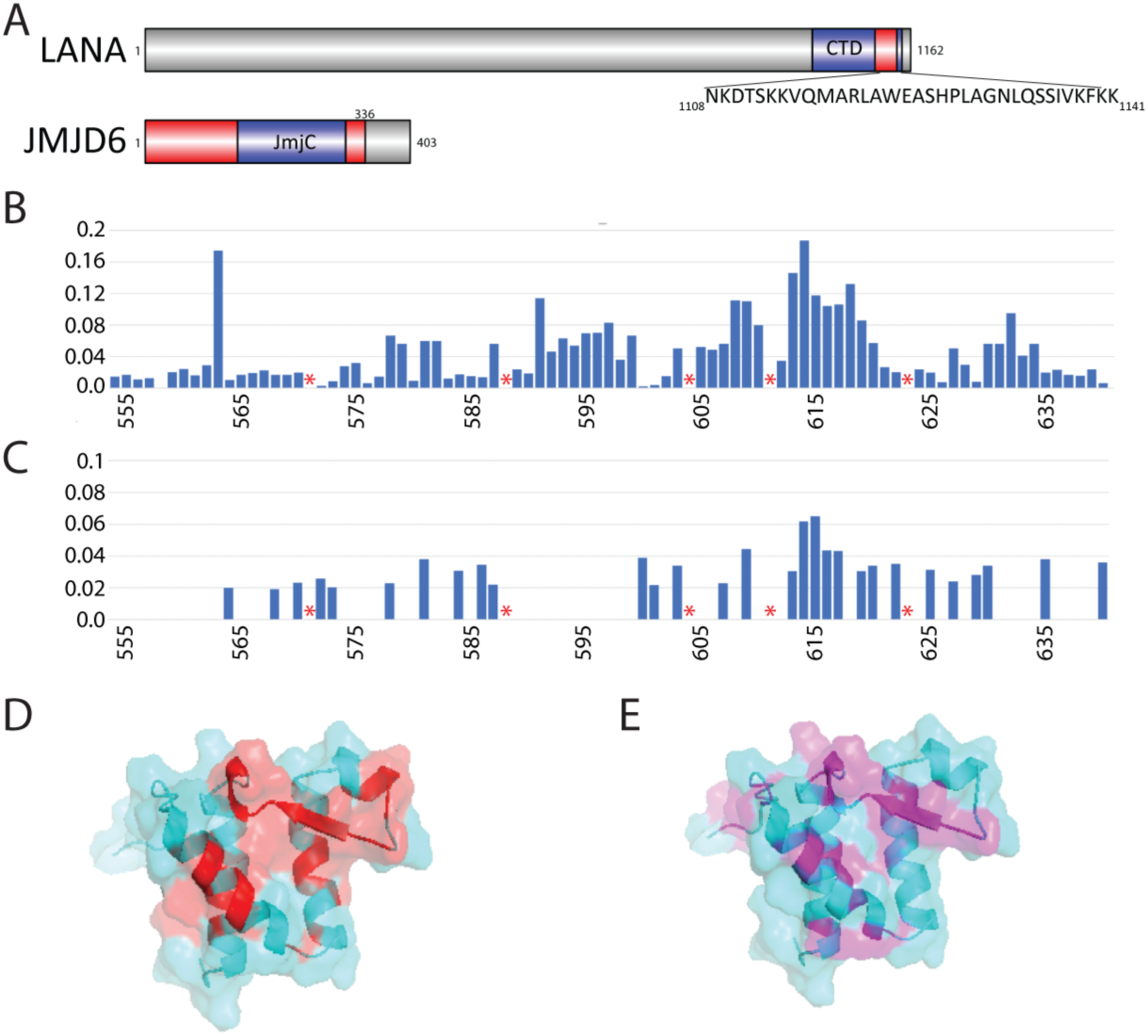
Comparison of chemical shift perturbations of BRD3 ET induced by LANA and JMJD6. (A) Constructs of LANA and JMJD6. Chemical shift perturbations of BRD3 ET domain relative to the bound (B) LANA and (C) JMJD6 calculated using Δδ(N,H)= ((δ_N_*6)^2^+(δ_H_)^2^)^0.5^ and plotted as a function of BRD3 ET sequence. Amino-acid residue positions marked with a red * correspond to Pro. The residues that were not assigned in the spectrum due to ambiguity of the assignment or missing cross peaks in the complex are included with Δδ(N,H)=0. (D) and (E) are the perturbations mapped onto the 3D structure of BRD3 ET and IN TP (IN TP is not displayed for ease of visualization) indicating the major interactions of the peptides are in the region of _613_EIEID_617_ in both the complexes in addition to those shown.

